# Platinum chemotherapy induces lymphangiogenesis in cancerous and healthy tissues that can be prevented with adjuvant anti-VEGFR3 therapy

**DOI:** 10.1101/781443

**Authors:** Alexandra R. Harris, Savieay Esparza, Mohammad S. Azimi, Robert Cornelison, Francesca N. Azar, Danielle C. Llaneza, Maura Belanger, Alexander Mathew, Svyatoslav Tkachenko, Matthew J. Perez, Claire Buchta Rosean, Raegan R. Bostic, R. Chase Cornelison, Kinsley M. Tate, Shayn M. Peirce-Cottler, Cherie Paquette, Anne Mills, Charles N. Landen, Jeff Saucerman, Patrick M. Dillon, Rebecca R. Pompano, Melanie A. Rutkowski, Jennifer M. Munson

## Abstract

Chemotherapy has been used to inhibit cancer growth for decades, but emerging evidence shows it can affect the tumor stroma unintentionally promoting cancer malignancy. After treatment of primary tumors, remaining drugs drain via lymphatics. Though all drugs interact with the lymphatics, we know little of their impact on them. Here, we show a previously unknown effect of platinums, a widely used class of chemotherapeutics, to directly induce systemic lymphangiogenesis and activation. These changes are dose-dependent, long-lasting, and occur in healthy and cancerous tissue in multiple mouse models of breast cancer. We saw similar effects in human ovarian and breast cancer patients whose treatment regimens included platinums. Carboplatin treatment of healthy mice prior to mammary tumor inoculation increases cancer metastasis as compared to no pre-treatment. These platinum-induced phenomena could be blocked by VEGFR3 inhibition. These findings have implications for cancer patients receiving platinums and may support the inclusion of anti-VEGFR3 therapy into treatment regimens or differential design of treatment regimens to alter these potential effects.

**Summary:** Platinum chemotherapy induces VEGFR3-dependent lymphangiogenesis, priming tissues for metastasis of breast cancer. Inhibition of VEGFR3 via antibody blockade can reverse these effects.

## Introduction

Over 650,000 cancer patients receive chemotherapy in the United States every year, with platinums, taxanes, and anthracyclines representing the most common classes of drugs (*1*). Unfortunately, many of these patients suffer recurrence. In breast cancer, the second most common cause of cancer death in women in the US, mortality results from metastasis rather than primary tumor growth (*2*). Ovarian cancer is similarly deadly due to dissemination of tumor rather than initial growth (*3*). Despite the centrality of metastasis to patient outcomes, it remains unclear why tumors that appear to be controlled or even cleared by initial chemotherapy later recur.

Chemotherapeutic drugs enter the tumor via blood vasculature and drain through the stroma toward peritumoral lymphatics. Similarly, tumor cells metastasize from carcinomas primarily via the lymphatics and downstream lymph nodes (*4*). Thus, the lymphatics represent a point of access for both tumor cells and chemotherapies to the rest of the body. Enlargement, sprouting, and proliferation of lymphatic vessels at both the primary tumor site (*5*, *6*) and metastatic sites (*7*) are associated with increased cancer growth, metastasis, and poor prognosis (*8*). Treatment of lymphangiogenic tumors with inhibitors of VEGF Receptor 3 (VEGFR3), which targets lymphatic endothelial cells (LECs), inhibits systemic metastasis (*9*, *10*).

Although chemotherapies pass through lymphatics, thus interacting with these gatekeeping vessels, little information is available on how chemotherapies affect them (*9*, *10*). Although one class of chemotherapy – taxanes – was shown to induce lymphangiogenesis in mice (*10*), here we examine how the other major classes of chemotherapy impact the lymphatics, with an emphasis on platinum chemotherapy. To our knowledge, no studies have been conducted that investigate the direct effects of chemotherapy on lymphatic endothelial cells. Platinum agents are some of the most commonly used chemotherapies in the United States and are currently administered to treat a variety of cancers (*11*). Thus, we aimed to understand how they impact lymphatic endothelial cell behavior and the implication of these changes on downstream cancer outcomes.

## Results

### Lymphatic endothelial cells respond to platinum chemotherapy

To assess if chemotherapeutics directly affect lymphatic endothelial cells, we treated monolayers of human LECs with low doses of docetaxel, doxorubicin, or carboplatin. Breakdown of LEC junctions is associated with vessel permeability, enhancing opportunities for tumor cells to intravasate (*12*). We observed dramatic morphological changes to LECs that included large gaps between cells in previously confluent monolayers after treatment with carboplatin (**Fig. 1A, B**), as well as other platinum agents, cisplatin and oxaliplatin (**Fig. S1A, B**), but not with doxorubicin or docetaxel (**Fig. S1C, D**). 95% of lymphatic endothelial cells exhibited junctional gaps with platinum treatment, compared to only 15% with vehicle treatment (**Fig. 1C**). Additionally, the severity of this junctional breakdown increased substantially, with platinum-treated LECs displaying junctional gaps nearly 7 times the size of vehicle-treated control LECs (**Fig. 1D**). Cellular adhesion molecules on LECs facilitate immune cell trafficking but are hijacked by tumor cells entering lymphatic vessels (*13*). LEC monolayers treated with carboplatin displayed higher numbers of ICAM1+ cells in hotspot regions compared to vehicle (**Fig. 1E,F**).

**Figure 1.**
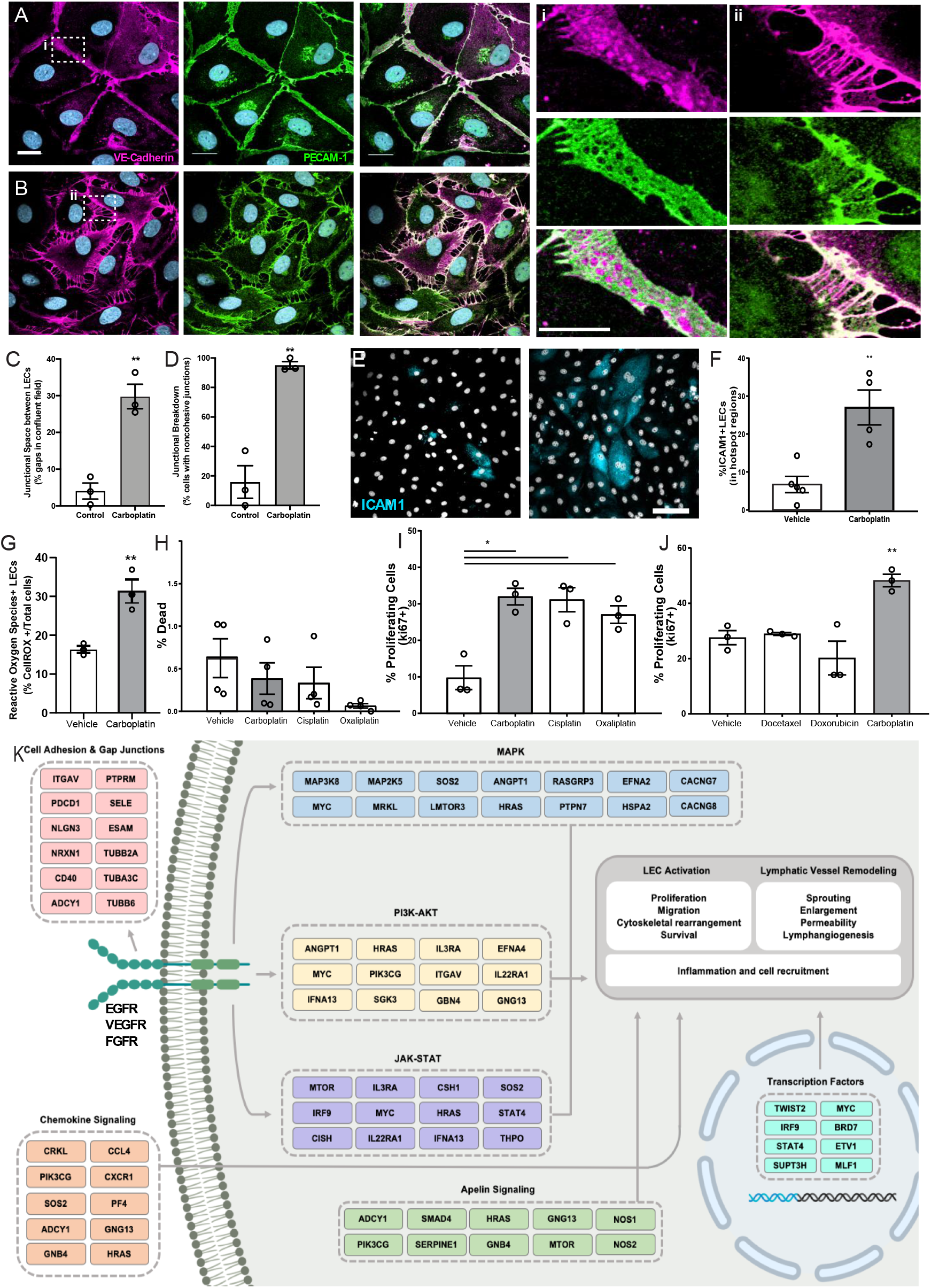
Lymphatic endothelial cell (LEC) monolayers are activated by carboplatin. (**A)** Untreated human LECs in monolayer culture; VE-cadherin (magenta) and PECAM-1 (green) with DAPI (gray-blue) for nuclei (scale bar=25μm) with (i) high magnification image cell-cell junction. **(B)** Human LEC monolayer treated with 1μM carboplatin with (ii) high magnification image of cell-cell junction. **(C)** Size of gaps between cells within a single field of view; listed as percentage of the field that is comprised of intracellular gap space. **(D)** Percentage of cells within fields of view that display non-cohesive junctions. **(E)** Representative images of LECs nuclei (DAPI, gray) and ICAM1 (cyan). Scale bar=250μm. **(F)** Percent ICAM1+ LECs in hotspot regions. **(G)** Percent ROS+ LECs per field. **(H)** Percent dead cells per field after 48h of platinum agent (1μM) assessed by amine-based fluorescent reactive dye. **(I)** Percent proliferating cells per field by KI67 positivity after 6h in culture with platinum agents (1 μM). **(I)** Percent proliferating cells per field by Ki67 positivity after 6h in culture with chemotherapeutic agents (1 μM). Each data set represents a minimum of three biological replicates. *p<0.05, **p<0.01. **(K)** Proposed mechanism generated from pathways enriched in LECs after *in vitro* treatment with carboplatin compared to vehicle-treated control. Gene members of each pathway were grouped reflecting their functional role in the cell.

Platinum chemotherapy induces cellular stress and can lead to production of reactive oxygen species (ROS). Further, ROS has been shown to disrupt tight junctions in endothelial cells and activate ICAM-1 and other cell adhesion molecules(*14*, *15*). Therefore, we examined ROS production in LECs treated with or without platinum. Carboplatin led to a significant 2-fold increase in the number of ROS+ LECs compared to control (**Fig. 1G**). ROS can have a dualistic role in cell signaling. In some contexts, ROS can trigger insurmountable oxidative stress that leads to cell death. Conversely, in other contexts, ROS can promote cell survival and even proliferation. To understand the potential implications of heightened ROS in LECs after chemotherapy, we chose to examine both cell death and proliferation in LECs treated with carboplatin. Interestingly, platinum treatment (24h) did not diminish viability (**Fig. 1H**); in fact, it induced a 3-fold increase in LEC proliferation, a necessary precursor for *in vivo* lymphangiogenesis (**Fig. 1I, S1E-G**). Platinums were the only class of chemotherapy tested that induced significant proliferation in LECs (**Fig.1I**). Together, these data suggest that platinum chemotherapy acts directly on LECs to induce phenotypes indicative of elements of lymphangiogenesis.

Gene set enrichment analysis of microarray data of LECs treated with the same doses of carboplatin reinforced these phenotypic changes, showing upregulation of pathways associated with proliferation, survival, and neovascularization (**Fig. 1J, Table S1**). Carboplatin induced activation of the pro-survival and proliferation pathways MAPK, JAK-STAT, PI3K/AKT, RAS, and HIF1-α signaling in LECs. Induction of these pathways, paired with increased expression of genes such as *MTOR*, *INOS*, *ANGPT1*, *CMYC*, *PI3K*, and others, may suggest platinums are activating growth factor signaling in LECs in response to cellular stress. Reverse phase protein array (RPPA) and pathway analysis indicated similar pathway upregulation, pointing to signaling via VEGFR, FGFR, and EGFR families **(Table S2**). Other enriched pathways included those governing cellular adhesion molecules, GAP junctions, and chemokine signaling, all important in LEC activation. Pathway enrichment analysis of the top upregulated microRNAs (9 in total, >2-fold increase in expression) showed similar pathway activation (**Table S3**).

### Carboplatin induces dose-dependent and sustained lymphangiogenesis in healthy murine, rat, and human tissues

Building on these *in vitro* data, we tested lymphangiogenesis in physiologically relevant models. First, we used *ex vivo* rat mesentery tissue (*16*) to analyze impact of treatment on lymphatics within intact vascular networks (**Fig. 2A**). Carboplatin significantly increased sprouting of lymphatic vessels by 3.5-fold (**Fig. 2B-D**) but had no detectable effect on blood vessels (**Fig. S2A-C**).

**Figure 2:**
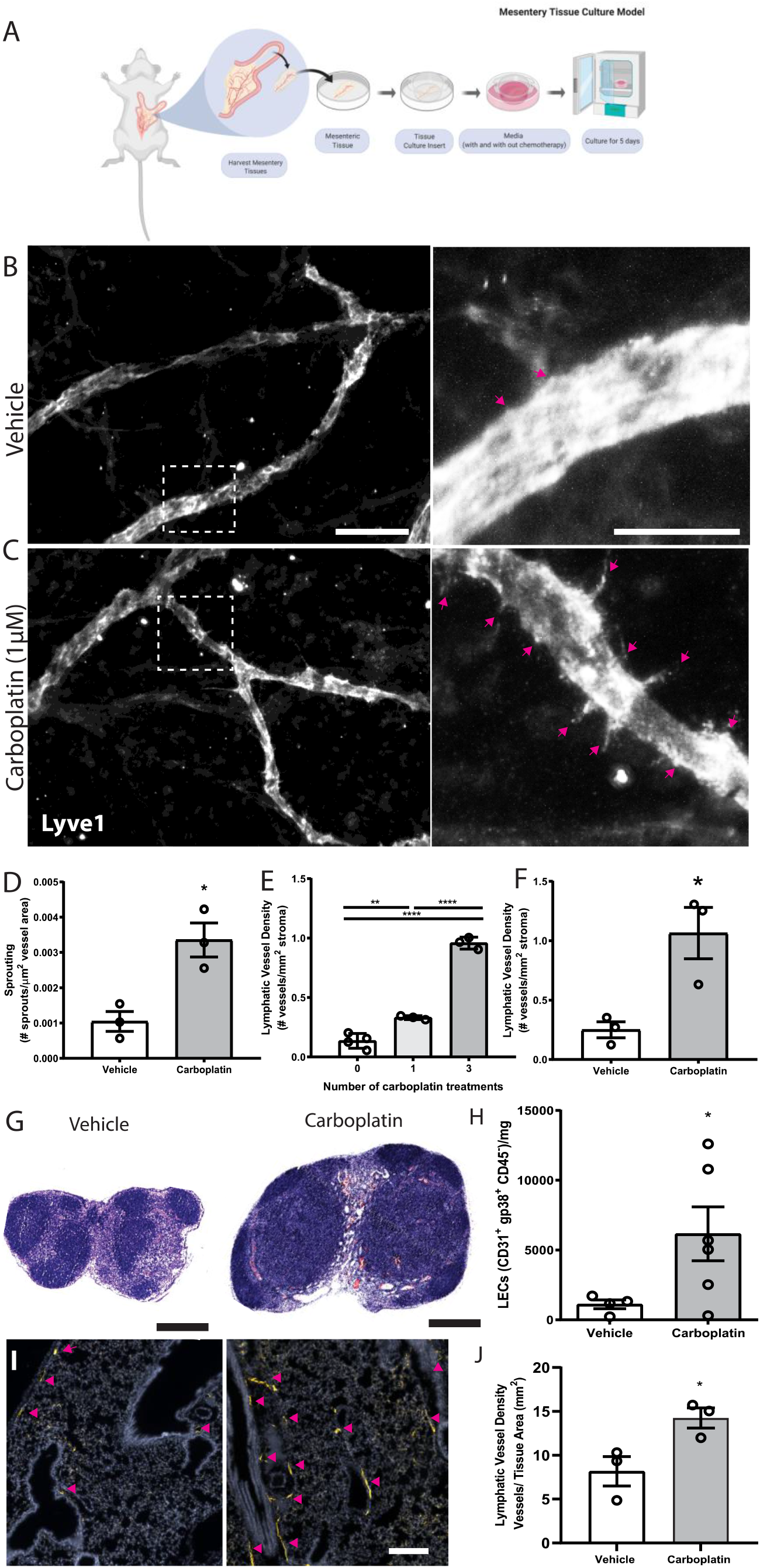
Carboplatin induces lymphangiogenesis in healthy tissues. **(A)** Schematic of rat mesentery culture model. **(B)** Vehicle-treated lymphatic vessels from mesentery cultures stained with LYVE-1 (grey). (**i**) High magnification image of boxed area in **B**. **(C)** Carboplatin-treated lymphatic vessels from mesentery cultures stained with LYVE-1 (grey).(**ii**) High magnification image of boxed area in **B**. Scale bar=100μm. **(D)** Number of sprouts per lymphatic vessel area. n=3. **(E)** Lymphatic vessel density (podoplanin+ vessels per mm^2^ stroma) in whole mammary fat pads of healthy mice treated with systemic carboplatin (8 mg/kg/dose) or vehicle by IV. n=3-4. **(F)** Lymphatic vessel density measured in mammary fat pads of healthy mice 2 months after treatment with 3 doses of carboplatin or vehicle, n=3. (**G)** Lymph nodes from healthy, tumor-naïve mice treated with vehicle and stained H&E. (**H)** LEC number in vehicle-treated and carboplatin-treated lymph nodes *in vivo.* **(I)** Representative images of lungs from mice treated with 3 doses of vehicle (*left*) and carboplatin (*right*). Lymphatic vessels noted by arrowheads. **(J)** Lymphatic vessel density in stromal tissue of lungs of mice pre-treated with carboplatin. *p<0.05, **p<0.01, ****p<0.001.

The importance of lymphatics in mammary carcinoma progression is well-established. We treated healthy, non-tumor-bearing female mice with 0-3 doses of carboplatin (8 mg/kg/dose) to examine lymphangiogenesis in naïve mammary fat pads (MFP) (**Fig. 2E**), histologically quantifying lymphatic vessel density (LVD), area, and perimeter (**Fig. S3**). Carboplatin treatment resulted in significant dose-dependent increases in LVD in the MFP stroma of both Balb/c and SCID mice (**Fig. 2E, Fig. S3G**), but no significant increase in vessel area or perimeter (**Fig. S3H,I).** LVD remained elevated 8 weeks after final treatment with carboplatin comparable to that at day 3 (**Fig. 2F**).

The observed lymphangiogenic effect of carboplatin on healthy mesentery suggests its lymphangiogenic effect is not restricted to mammary tissue. As lymph nodes are a frequent site of cancer spread away from the primary tumor, we also examined lymph node lymphangiogenesis in healthy mice treated with carboplatin chemotherapy. Carboplatin treatment resulted in larger inguinal LNs (**Fig. 2G)**, and showed a significant increase in LEC number (**Fig. 2H**).

In addition to lymph nodes, we were also curious to examine the effect of platinum on lungs. Lungs are one of the most common metastatic sites in cancer and lymphangiogenesis in lung tissue can enhance metastatic seeding by creating more exit points for disseminating cancer cells (*17*). Indeed, platinum treatment of healthy mice stimulated lung lymphangiogenesis, leading to a nearly 2-fold increase in lung LVD after carboplatin (**Fig. 2I, J**).

### Chemotherapy regimens that include platinum agents are associated with higher LVD in patients

Platinum chemotherapy is standard of care in high-grade serous ovarian cancer along with cytoreductive surgery. Because the omentum is the most frequent site of ovarian cancer metastasis, it is routinely removed during surgery, giving us access to histologically normal tissues. We analyzed lymphatics in omentum from patients treated with neoadjuvant carboplatin and docetaxel chemotherapy prior to surgery (**Table S4**). All omentum samples were pathologist-identified as uninvolved and ostensibly healthy. Histologically normal omentum treated with carboplatin had 12-fold higher LVD compared to that of untreated patients (**Fig. 3A,B**). Though we could not feasibly procure patient tissues treated with only platinum due to carbotaxol standard of care, our previous findings in animal models have demonstrated that taxanes require tumor to promote lymphangiogenesis (*10*). Thus, the higher LVD observed here are likely attributable to carboplatin, concordant with our results *in vitro* and in rodents.

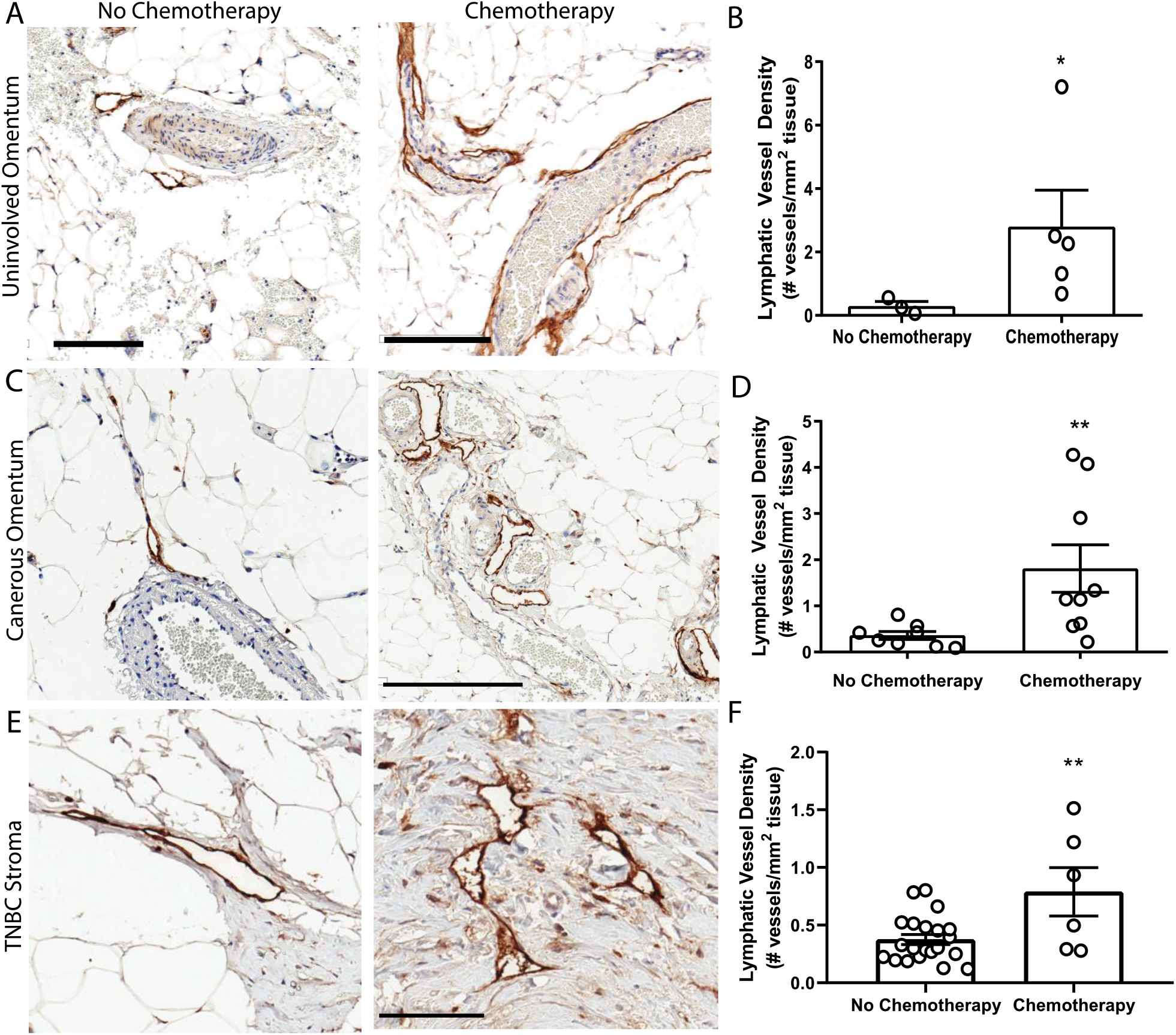
Platinum chemotherapy is associated with higher LVD in human cancer patients. **(A)** Representative images of lymphatic vessels in histologically benign omentum from patients treated with no chemotherapy (top) or neoadjuvant carboplatin combination chemotherapy (bottom) (*see Supplemental Table 4*) podoplanin (brown) and hematoxylin (blue). (**B**) Lymphatic vessel density in patient samples. Scale bar = 200 microns. N=8 patients. **(C)** Representative images of tissues from ovarian cancer patients treated with no chemotherapy (*left*) or neoadjuvant carboplatin and paclitaxel (*right*) in cancerous omentum with podoplanin(brown) and hematoxylin(blue). (**D**) Quantified lymphatic vessel density (N= 17). **(E)** Representative images of tissues from triple negative breast cancer patients treated with no chemotherapy (*left*) or neoadjuvant platinum and taxane (*right*) with podoplanin(brown) and hematoxylin(blue). (**F**) Lymphatic vessel density (N=27) *p<0.05, **p<0.01, scale bar=200μm.

In addition to our analysis of pathologically normal tissues, we quantified LVD in cancerous omental tissue from 17 ovarian cancer patients with or without carbotaxol chemotherapy prior to surgery (**Fig. 3A, Table S5**). We again detected a significantly higher LVD in patients that received neoadjuvant platinums (**Fig. 3B**). In addition to gynecologic malignancies, platinums are routinely used in the clinical management of triple negative breast cancer (TNBC) often as a neoadjuvant treatment prior to surgery. Similarly, in primary tumor stromal tissue from 27 TNBC patients (**Fig. 3C, Table S6**), there was a significantly higher LVD in patients treated with platinums prior to surgical resection (**Fig. 3D**)

### Platinum chemotherapy induces lymphangiogenesis in breast tumor stroma

To experimentally investigate causal effects of platinum on tumor lymphangiogenesis, we employed a series of preclinical breast cancer models: orthotopic 4T1 syngeneic tumors (immune-competent); orthotopic MDAMB231 xenograft (immune-compromised); or the inducible autochthonous mammary tumor model, L-Stop-L-K-*KRas^G12D^p53^flx/flx^*-L-Stop-L-*Myristoylated p110α*-GFP (immune-competent), which more closely mimics the malignant transformation and progression in humans (*18*). We chose breast cancer as a model for these studies because breast cancer preferentially metastasizes via lymphatics(*4*), unlike ovarian cancers that can spread through multiple mechanisms. Mice received carboplatin when tumors were palpable and were treated in the same dosing and schedule as previously discussed. There were no differences in tumor size observed (**Fig. S4A**). Histological analysis (**Fig. 4A-D**, **Fig. S4B,C**) showed significantly increased LVD after three treatments of carboplatin (**Fig. 3E**). Unlike in naïve MFP, lymphatic vessel enlargement was present, with lymphatic vessel area and perimeter significantly increased in immunocompetent mice (**Fig. 3F, S4D**).

**Figure 4:**
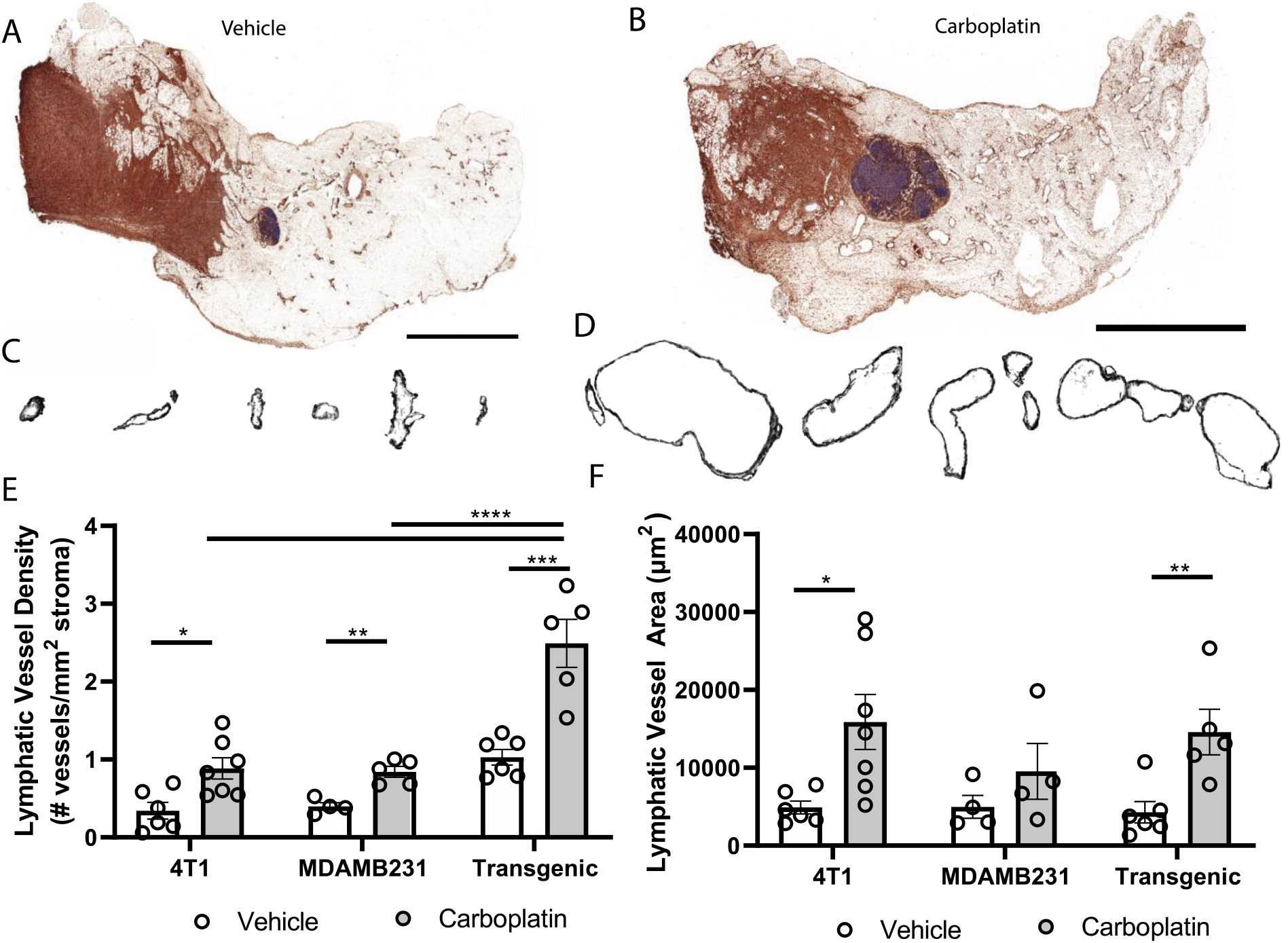
Platinum agents induce lymphangiogenesis in the tumor stroma. **(A)** Representative tumor-bearing mammary fat pad with podoplanin (brown) and hematoxylin (blue) from vehicle-treated transgenic Kras^G120D^p53^fl/fl^p110a^myr^ mice. **(B**) Representative whole tumor-bearing mammary fat pad from carboplatin-treated transgenic mice. (**C**) Representative individual lymphatic vessel cross-sections from **A**. (**D**) Isolated lymphatic vessels from **B**. (**E**) Lymphatic vessel density (lymphatic vessels per stromal area) in tumor-bearing mammary fat pads in 3 orthotopic mouse models of breast cancer (syngeneic 4T1, transgenic Kras^G120D^p53^fl/fl^p110a^myr^, and xenografted MDA-MB-231). (**F**) Average area of individual lymphatic vessels (μm^2^) in mouse models of breast cancer as described in **I**. *p<0.05, **p<0.01, ***p<0.001, ****p<0.0001, n≥4/cohort.

### Systemic carboplatin pre-treatment leads to increased lymph node and lung metastasis in murine breast cancer

Enlargement and remodeling of lymphatic vessels is known to increase tumor cell dissemination (*6*, *8*, *19*), with LVD and tumor spread correlating in murine and human cancers (*20*–*22*). Using a tissue engineered model of the breast cancer microenvironment, we also saw that addition of carboplatin in the presence of lymphatic endothelial cells would increase tumor cell invasion towards lymphatics (**Fig. S5A,B**). We hypothesized that priming of healthy tissues with carboplatin creates a hospitable niche that will later promote tumor cell invasion and metastasis once the cancer disease process has started.

We treated healthy, tumor-naïve mice with systemic carboplatin to induce lymphangiogenesis (**Fig. 5A**), followed by orthotopic implantation of 4T1 tumor cells 1 week later. Since carboplatin was allowed to clear completely prior to cancer initiation, tumors were never exposed to the drug and thus there was no significant difference in tumor size at endpoints, as expected (**Fig. S5C**). As we previously observed that platinum increases LEC numbers in the lymph node and lung, we assessed metastasis to those organs, which are common sites of cancer spread. Tumor cell metastasis to the tumor-draining inguinal lymph node (TDLN), considered a first step in tumor cell dissemination (*4*), significantly increased in platinum pre-treated TDLNs compared to control (**Fig. 5B-D**). These data suggest that the lymphatics activated and remodeled by platinums are functionally capable of promoting metastasis. We then evaluated whether platinum-priming could contribute to distant metastases, e.g. to lung. 21 days after implantation, 100% of carboplatin pre-treated mice showed gross lung metastases, compared to 50% of the vehicle. Microscopic examination of lungs showed significantly increased numbers of foci (**Fig. 5E,F**) that were significantly larger with pretreatment (**Fig. 5G**). Therefore, we find that systemic pre-treatment with carboplatin increases LN and lung metastasis.

**Figure 5:**
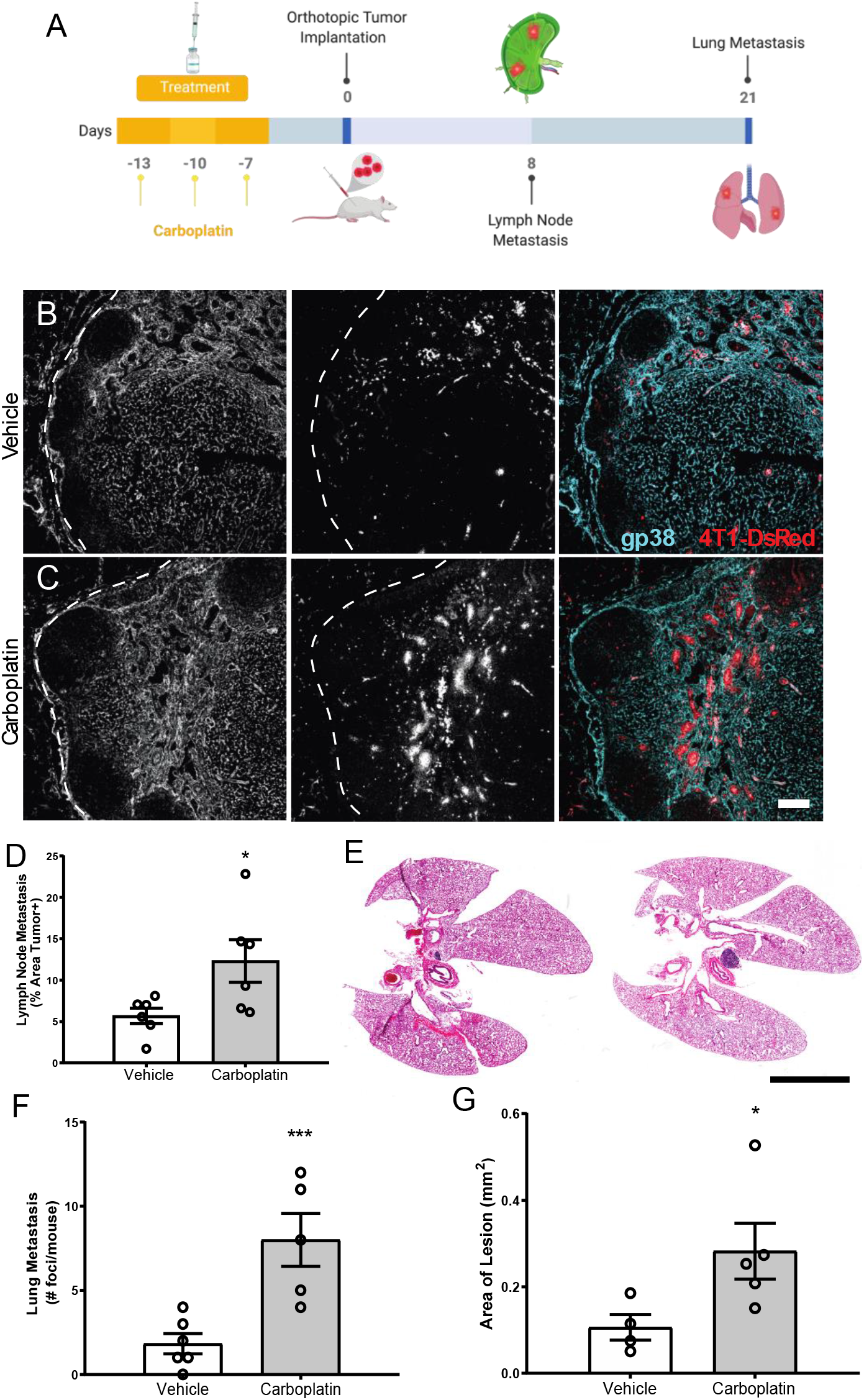
Systemic pre-treatment with carboplatin increases metastatic spread of 4T1 breast tumors. **(A)** Experimental timeline for carboplatin pretreatment (8 mg/kg × 3 or vehicle) followed by 4T1 tumor implant and subsequent tissue harvest. **(B)** Representative images of 4T1 cells (red) in vehicle pretreated tumor-draining inguinal lymph nodes (TDLN) counterstained with podoplanin (cyan). **(C)** Representative images from carboplatin pretreated TDLNs. Scale bar=100μm. **(D)** Percentage of TDLN area covered by tumor cells as assessed by image thresholding in ImageJ. **(E)** Cross-section of lungs from vehicle (left) and carboplatin (right) -primed mice by H&E. Scale bar=1mm (**F**) Number of metastatic foci in lung per mouse. **(G)** Area of macroscopic metastatic lesions in lung. *p<0.05, ***p<0.005 as analyzed by individual *t* test, n≥4.

### VEGFR3 blockade mitigates carboplatin-induced lymphangiogenesis and metastasis

VEGFR3 blockade reduces metastatic spread and lymphangiogenesis in a number of tumor models, as the VEGFC:VEGFR3 signaling pathway is one of the quintessential drivers of LEC proliferation and lymphatic expansion (*6*). Based on the literature implicating VEGFR3 in metastatic spread in murine breast cancer models, its specificity for lymphatics, as well as the potential connection to VEGFR3 as seen in our gene set enrichment analysis (Fig. 1J), we hypothesized that blockade of VEGFR3 would abrogate the effects of carboplatin on lymphatics. Thus, we examined outcomes in the presence of anti-VEGFR3 inhibition.

We used MAZ51 (3-(4-Dimethylamino-naphthalen-1-ylmethylene)-1,3-dyhydro-indol-2-one), a specific small molecule inhibitor of VEGFR3, on monolayers of LECs *in vitro*. MAZ51 had little effect on LEC proliferation alone, but significantly suppressed carboplatin-induced LEC proliferation (**Fig. 6A**) and junctional disruption (**Fig. S6A)** to resemble untreated LECs (**Fig. 1**). Combination treatment with carboplatin and MAZ51 only modestly increased LEC death from less than 1% to ~3% (**Fig. S6B,C**).

**Figure 6:**
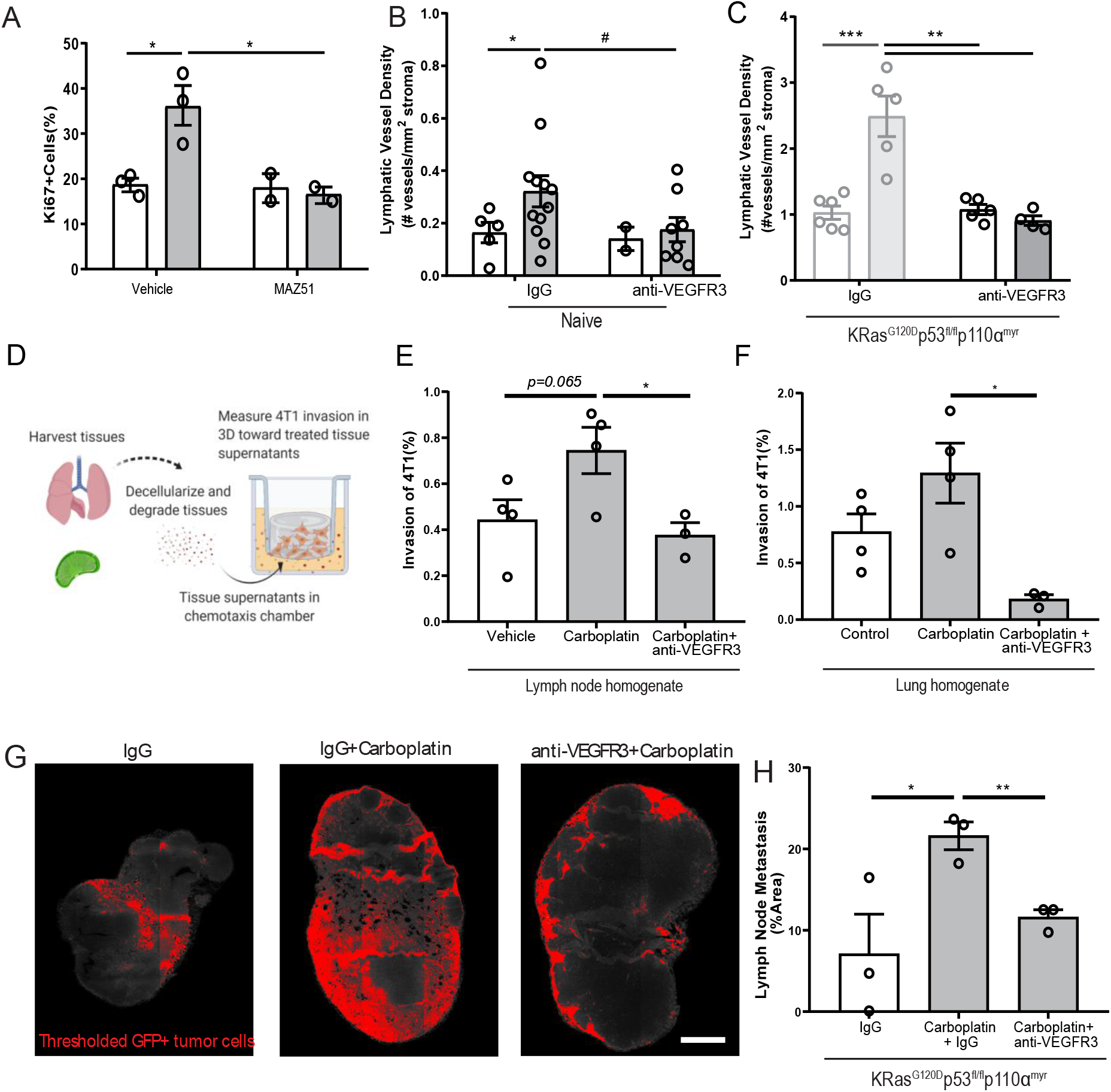
Blockade of VEGFR3 inhibits carboplatin-induced lymphangiogenesis and metastasis. **(A)** Percent proliferating cells per field by Ki67+ staining of human LEC monolayers treated *in vitro* with carboplatin (1 μM), VEGFR3 inhibitor MAZ51 (1 μM), or both for 6h. **(B)** Lymphatic vessel density in naïve, tumor-free fat pads from mice treated with vehicle/carboplatin and anti-VEGFR3/ IgG. **(C)** Lymphatic vessel density (lymphatic vessels per stromal area) in tumor-bearing fat pad of KRas^G120D^p53^fl/fl^p110α^myr^ mice treated vehicle/carboplatin and anti-VEGFR3/IgG. Grayed bars represent data presented in previous figure but included here as point of reference. **(D)** Experimental schematic of chemotaxis assay using digested and decellularized *in vivo*-treated tissues. Healthy, tumor-free mice were treated in vivo with carboplatin (8 mg/kg × 3 or vehicle) and/or anti-VEGFR3 antibody (or control IgG antibody). Lymph nodes and lungs were harvested 3 days following final treatment, digested and decellularized, and used in a 3D *in vitro* chemotaxis assay for 4T1 cells. **(E)** Invasion of 4T1 tumor cells towards digested and decellularized lymph node **(E)** and lung **(F)** from treated mice. **(G)** Representative thresholded images and **(H)** quantification GFP+ tumor cells in whole inguinal lymph nodes from KRas^G120D^p53^fl/fl^p110α^myr^ mice treated with vehicle/carboplatin and anti-VEGFR3/IgG.

Inhibition of VEGFR3 fully attenuated LEC proliferation caused by platinums. We speculated that platinum-induced lymphangiogenesis in tumor-free tissues was also dependent on VEGFR3 signaling. Indeed, we observed the same trend in the mammary fat pads of mice treated with a clinically relevant VEGFR3 blocking antibody, where VEGFR3 inhibition reduced LVD to baseline levels (**Fig. 6B**). In tumor-bearing tissue from our breast cancer animal models, we could reverse carboplatin-mediated increases in LVD in both 4T1 (**Fig. S7A**) and KRas^G120D^p53^fl/fl^p110α^myr^ **(Fig. 6C)** tumor-bearing fat pads. Similarly, blockade significantly reduced tumor-associated vessel area (**Fig. S7B,D**) and perimeter (**Fig. S7C,E)**. Importantly, anti-VEGFR3 co-treatment fully reversed the platinum-induced lymphangiogenesis in murine lungs of healthy mice treated with platinums *in vivo* **(Fig. S7F).** These data suggest the changes to lymphatics by platinum chemotherapy is mediated by VEGFR3 and its inhibition can reverse these phenomena.

With platinums, we observed increased activation (Fig. 1) and proliferation (Fig. 1, 2) of lymphatics in otherwise healthy tissues throughout the body. The lymphatic endothelium secretes a number of chemotactic factors that can promote pre-metastatic niche formation and subsequent tumor cell invasion. To understand whether factors secreted by lymphatics in the tissue milieu could enhance tumor cell chemotaxis, we treated healthy, tumor-free mice with carboplatin and anti-VEGFR3 antibody *in vivo* and harvested their lymph nodes and lungs three days after the final treatment (**Fig. 6C-E**). These tissues were digested and decellularized (**Fig. 6D**); their supernatants were then used in a chemotaxis chamber to measure invasion of murine breast cancer cells through a 3D matrix. Tumor cells invaded more to carboplatin-treated lymph node (**Fig. 6E**) and lung (**Fig. 6F**) digested and decellularized tissues. Further, tissue supernatants from mice that were also treated with anti-VEGFR3 antibody *in vivo* showed significantly attenuated platinum-induced invasion of 4T1 cells toward both tissue homogenates. These data indicate that the activating effects of platinum on lymphatics that promote tumor cell invasion are mediated through VEGFR3.

Ultimately, metastasis reduction *in vivo* is desired. While carboplatin treatment increased lymph node metastasis by nearly 3-fold (**Fig. 6G,H**), alternating carboplatin with anti-VEGFR3 therapy (**Fig. S7G,H**) resulted in reversion of metastatic spread induced by carboplatin in KRas^G120D^p53^fl/fl^p110α^myr^ mice (**Fig. 6G,H**). Taken together, these data indicate that carboplatin-associated lymphangiogenesis occurs via a VEGFR3-dependent mechanism. VEGFR3 inhibition could directly revert the phenotypic effects of carboplatin seen *in vitro*, *ex vivo*, and *in vivo,* thereby explaining its therapeutic success.

## Discussion

Here we see that platinum agents, and most specifically carboplatin, increase lymphatic expansion and proliferation across a number of models. The phenotypic changes we observed with platinums *in vitro* were indicative of changes *in vivo*. Though the pathway we ultimately targeted to reduce the phenotype, VEGFR3, is non-novel for lymphangiogenesis, the connection to carboplatin (and to a lesser degree, platinum agents in general), is. Accordingly, we observed increases in LEC numbers and LVD *in vitro*, *ex vivo*, *in vivo*, and in patient tissues with platinums. It is counterintuitive that chemotherapies would induce expansion of a cellular population and promote tumor progression, however, in fibroblasts, induction of stress responses by DNA-damaging chemotherapies can result in similar activation and compensatory proliferation (*27*, *28*). Platinums have been shown to increase ROS(*34*), and ROS can lead to enhanced proliferation, survival, and inflammatory pathways(*35*), all of which we observed in our LECs after treatment. Interestingly, ROS, of which we see an increase, can induce oxidation of growth factor receptors and downstream phosphatases to enhance growth factor receptor signaling(*36*). Therefore, though we know that VEGFR3 blockade can reduce the pro-lymphangiogenic effects that we see, the full mechanism requires further exploration to determine the link between platinum agents, ROS, and lymphangiogenesis across physiological contexts.

Platinum-induced lymphangiogenesis occurred in multiple tissues, such as mammary tissues, lymph nodes, lungs, and connective tissues, and was sustained long-term. Permanent increases and remodeling to lymphatics not only at the primary tumor site but also in distant tissues may prime these tissues by encouraging formation of the pre-metastatic niche, creating more potential escape routes for disseminated tumor cells. In fact, this is the case in cancers such as colorectal and breast cancer, where increased LVD in the liver and lungs, respectively, has been correlated to increased instance of metastasis in those tissues (*38*); indeed, we observed increased LVD in the lungs of platinum-treated mice that was VEGFR3-dependent. This is in direct contrast to lymphatic effects seen with other chemotherapeutic agents like taxanes that require the presence of a tumor to enact these changes and occur only in the local tumor microenvironment (*9*, *10*). Differences in these mechanisms may arise from the inherent differences in drug mechanism of action between taxanes, which are not DNA damaging, and platinums, which act directly on DNA. As patients are exposed to chemotherapies on a systemic level, platinum-induced lymphangiogenesis could conceivably occur in distant tissues posing an increased risk for lymphatic metastasis to these other sites.

All therapies, anti-cancer or otherwise, eventually drain via our lymphatic system, but the majority of research has focused on the impacts of drugs on the blood vasculature, largely ignoring the impact on this endothelium (*42*). To our knowledge, this is the first study that characterizes the effects of chemotherapy on lymphatics in healthy, cancer-free in vitro, ex vivo, and in vivo models. Indeed, more studies are emerging that suggest chemotherapy may paradoxically counteract its own efficacy through action on either the tumor cells or its associated stroma. Chemotherapies have myriad off-target effects, including enriching for cancer stem cell populations (*46*), promoting invasiveness (*47*), and inducing epithelial-to-mesenchymal transition (EMT) to encourage drug resistance (*48*). In this study, we examined the physiological implications of platinum-induced lymphangiogenesis in the context of metastasis, illustrating potential deleterious off-target effects of a commonly used therapy. However, some off-target effects of chemotherapy can also be desirable. We did not investigate the impact of platinum-induced lymphangiogenesis on tumor immunity, which may offer positive benefits by increased immune cell migration to the tumor and metastatic sites. Platinums and other chemotherapies can generate neoantigens and activate T cells for enhanced immunological response. Activated lymphatics can assist in immune cell trafficking to the tumor site. Through this lens, it is possible that the presence of increased tolerogenic LECs and normalized vasculature after VEGFR3 therapy could interfere with those benefits (*30*). Further investigation is needed into the physiologic implications of these lymphovascular changes in patients. Certainly, platinum chemotherapy has demonstrated potent cytoreductive efficacy over decades of clinical use. However, clinical decision-making cannot be truly informed until more studies examine how these agents affect normal tissues.

Our studies describe a previously unknown pro-lymphangiogenic action of platinum chemotherapeutics. In our pre-treatment animal models, we found that this led to enhanced metastasis. Platinum-induced lymphatic proliferation and expansion were dependent on VEGFR3 and, therefore, supplementing chemotherapeutic regimens with anti-VEGFR3 was sufficient to prevent them. VEGFR3 inhibition has been suggested as a potential anti-metastatic treatment option in human cancers since the discovery of this pathway as a leading driver of lymphangiogenesis. It has shown successful anti-metastatic benefits in multiple murine models of cancer (*5*, *19*, *49*–*52*) but had mixed results in early phase clinical trial. Our data indicate it may be most advantageous not as a single agent but instead in combination with a platinum chemotherapy as a preventative measure against countertherapeutic VEGFR3-dependent lymphangiogenesis. Patients with cancers that spread preferentially through the lymphatic system, such as carcinomas, that are at an early stage would likely receive the most benefit from this pairing. Conversely, patients in advanced stages already showing rampant hematogenous disease would likely not show added benefit from this approach; this may also offer an explanation for the subpar anticancer performance by anti-VEGFR3 in early-phase clinical testing. Clearly, more pre-clinical testing of this pairing is warranted to fully understand its clinical potential.

We believe that these findings highlight our incomplete understanding of chemotherapeutic action in the tissue stroma. Our data here suggest that informed pairing of chemotherapy with targeted therapies to the tumor microenvironment may improve overall efficacy of these valuable treatments. Specifically, we believe our findings may renew interest in the clinical potential of anti-VEGFR3 therapy, such as IMC-3C5 (*53*), which is well-tolerated, yet stalled in clinical trials, and may show promising results when used in combination with specific chemotherapy within the proper disease context.

## Materials and Methods

### Cell culture

Human lymphatic endothelial cells (HMVEC-dLy, Lonza) were cultured in Endothelial Cell Growth Medium (EBM-2 basal media, Lonza) supplemented with recommended growth supplement kit (EGM-2MV BulletKit, Lonza). Mouse mammary carcinoma cell line 4T1-luc-red (generously given by the Cross laboratory at University of Virginia) originated from ATCC and were acquired from Perkin-Elmer (BW124087V) after lentiviral transduction of Red-FLuc luciferase gene. 4T1 cells were cultured in RPMI medium supplemented with 10% FBS. MDA-MB-231 were acquired from the ATCC and cultured in Dulbecco’s modified Eagle’s medium (DMEM, Gibco) and supplemented with 10% fetal bovine serum (FBS). All cell lines were grown sterilely in humidified atmosphere of 5% CO2 and 95% oxygen at 37°C. Cell lines were tested for mycoplasma and all experiments were completed afterwards.

### *In vitro* drug treatment, immunocytochemistry, and live/dead assays

LECs were cultured on glass coverslips in complete media as described above; the LEC monolayer was treated with 1 μM carboplatin, cisplatin, oxaliplatin, docetaxel, doxorubicin, MAZ51(*54*), or appropriate solvent control (referred to as ‘vehicle’) for 6 hours (phenotypic studies) or 48 hours (live/dead analysis). After treatment, coverslips were fixed with 4% paraformaldehyde (PFA) for 30 minutes at room temperature and underwent immunofluorescent staining with Ki67 to assess proliferation (Millipore, Cat. #AB9260); ICAM1 to assess cellular adhesion molecule expression (Abcam, Cat. #AB2213); VE-Cadherin (Abcam, Cat. #AB33168) and CD31 (R&D systems, Cat. #AF806) to assess cellular junctions. For live/dead analysis, amine-based fixable live/dead solutions (Life Technologies, Cat. #L23101) were added to cell media of living LECs after 48 hours of drug treatment and five random images were taken of each well; technical replicates were averaged to yield one biological replicate. For VEGF-C measurements, LECs were treated with 1μM carboplatin for 6h and then media was collected and cells were lysed. VEGF-C was assessed using the R&D Quantikine ELISA kit. Each quantification was performed with a minimum of 3 biological replicates.

### CellROX Reactive Oxygen Species (ROS) Assay

Relative ROS generation was measured using CellROX Green Reagent (Thermo Fisher). This is a DNA-binding probe that fluoresces bright green upon oxidation by ROS. The assay was conducted as recommended by the supplier. In brief, LECs were cultured in complete media as described above; the LEC monolayer was treated with 1 μM carboplatin, MAZ51 (50), 100 uM Menadione K3 (Sigma, Cat.#47775) as a positive inducer of ROS(*55*), 1mM N-Acetylcysteine (Sigma, Cat.# A9165) as a negative control per manufacturer’s recommendation, or appropriate solvent control (referred to as ‘vehicle’) for 6 hours. After treatment, cells were washed with PBS and incubated with 2.5 μM CellROX Green Reagent and 1.62 uM Hoescht 33342 in the dark for 30 minutes at 37°C. Afterwards samples were washed once with PBS before measuring intracellular levels of ROS with a Zeiss Axio Observer fluorescent microscope. Five random images were taken of each well using the DAPI and FITC filter. Quantification was done using ImageJ, to count the number of cellROX green positive cells with signal present in both the nucleus and mitochondria being considered. To find CellROX+ percent of cells, number of CellROX+ cells was divided by total nuclei per area. For CellROX ROS analysis, means taken from each treatment triplicate were considered a biological replicate for statistical analysis.

### Microarray and Gene Set Enrichment Analysis

Human LECs were treated *in vitro* with 1 μM carboplatin for 4 hours. Cells were lysed with RLT buffer and RNA was isolated using the QIAGEN RNeasy Kit (Cat #74104). Microarray was performed by the UVA DNA Sciences core using the Human Affymetrix GeneChip Array ([HuGene-2_1-st] Affymetrix Human Gene 2.1 ST Array). Gene lists were generated from microarray data corresponding to genes >1.25 LogFC and <0.75 logFC were used in downstream analyses. Differential targets identified were further subjected to gene set enrichment analysis performed using the ShinyGO enrichment tool using the STRING api. The STRING database (Search Tool for the Retrieval of Interacting Genes/Proteins) is a tool for looking at functional associations between different proteins(*56*). Protein-protein interactions are mapped across several curated databases and functional association is mapped in interaction networks. Gene interactions are then mapped to the KEGG pathway annotations with the adjusted p values representing pathway enrichment analysis from KEGG Release 86.1. MicroRNAs upregulated by >2 LogFC after treatment were subjected to enrichment analysis by DIANA mirPath v.3 from KEGG pathways significantly enriched with the union of target genes(*57*).

### Reverse-phase protein array and pathway analysis

Human LECs were treated *in vitro* with 1 μM carboplatin for 6 hours. Cells were washed and lysed using RIPA buffer with protease and phosphatase inhibitors. RPPA was performed by the MD Anderson Functional Proteomics Core Facility as previously published(*58*). To assess which proteins reacted to the treatment, ratios of brightness were calculated and the distribution was assessed. Outlier proteins, for which the ratio was more than 2 standard deviations from the mean of the ratio distribution, were taken to be most affected by the treatment. Pathway enrichment analysis for these proteins was performed using ConsensusPathDB (*59*) online tool and pathways enriched with statistical significance of unadjusted p-value of 0.01 were retained for further analysis.

### Harvest, treatment, and whole mounting of *ex vivo* rat mesentery tissues

Mesentery tissues were harvested from rats as previously described by Azimi, et al(*16*, *60*) (**Fig. 2A**). Briefly, mesenteric tissue was harvested from the small intestine of an adult Wistar rat and transferred into a culture dish. Tissues were arranged on permeable membranes of cell crown inserts and cultured in MEM with 20% FBS and 1% penicillin/streptomycin for five days as previously described with 1 μM carboplatin or vehicle control. Tissues were then removed from inserts, grossed, mounted to slides, and fixed with methanol to undergo immunofluorescent staining with PECAM-1/CD-31 (BD Biosciences, Cat. #555026) for blood vessels and LYVE-1 (AngioBio Co., Cat. #11-034) for lymphatic vessels. Images were taken of a minimum of five random areas of vessel remodeling for each well and quantified by counting the number of sprouts normalized to vessel area in each field.

### *In vivo* animal study design

All methods were performed in accordance with relevant guidelines and regulations and approved by the University of Virginia Institutional Animal Care and Use Committee.

#### Studies in healthy, tumor-naive animals

6-week-old female Balb/c mice were treated via tail vein injection of carboplatin unless otherwise noted. Mice underwent three total treatments with 8 mg/kg carboplatin or vehicle control (saline); each treatment was three days apart and mice were euthanized CO_2_ inhalation three days following the final treatment. Naïve mammary fat pads and axillary lymph nodes were harvested; fat pads were post-fixed in 4% PFA for 24 hours, dehydrated, paraffin-embedded, and sectioned at 7-micron thickness to undergo immunohistrochemical staining (see Immunohistochemistry). Lungs were harvested, post-fixed in 4% PFA followed by 30% sucrose and embedded in OCT and sectioned at 12-micron thickness. Axillary lymph nodes were digested (dissociation as previously described(*61*)) and total LEC counts quantified (see Flow Cytometry).

#### Studies in 4T1 breast cancer model

4T1 mouse mammary carcinoma cells were cultured as described above. 4T1 cells were suspended in 3.3 mg/ml growth-factor reduced basement membrane extract (Cultrex) in phosphate-buffered saline (PBS) and orthotopically injected in a subareolar fashion into the fourth mammary fat pad of female balb/c mice. For most experiments, 50,000 4T1 cells were injected and mice treated with either 1 or 3 doses of 8 mg/kg carboplatin or vehicle by tail vein injection staggered by one day with anti-VEGFR3 antibody (100 μg per injection × 3 total injections, I.P., eBioscience (now ThermoFisher), Control IgG: rat monoclonal IgG2a kappa Isotype Control, Cat.#16-4321-85; Anti-VEGFR3: rat monoclonal IgG2a kappa to mouse VEGFR3 (AFL4): Cat. #16-5988-85) or IgG control antibody after tumors were just palpable. Mice were euthanized by CO_2_ inhalation once tumors reached desired endpoints as assessed by caliper measurement. Tumor-bearing and contralateral naïve fat pads containing inguinal lymph nodes were harvested and post-fixed (24 h for naïve tissues, 48 h for tumors) in 4% PFA and processed for histology as described above; tumor-draining axillary lymph nodes were dissociated for flow cytometric analysis for LEC number.

For **pre-treatment experiments**, naïve mice were treated with three rounds of carboplatin as previously described and drug was allowed one week to clear(*62*). 10,000 4T1 cells were then injected as described above and allowed to grow until desired size endpoints were reached. Tumor-bearing fat pads were harvested and processed as described above. Lungs were removed, washed in saline, and fixed in 4% PFA for 48 h where they were then embedded in OCT and cryosectioned to undergo histological staining to examine metastasis.

#### Studies in transgenic breast cancer model

L-Stop-L-*KRas^G12D^p53^flx/flx^*L-Stop-L-*Myristoylated p110α*–GFP^+^ mice on a C57BL/6 background were generously provided by Melanie Rutkowski, University of Virginia. Mammary tumors were initiated by intraductal injection of adenovirus-Cre in these mice as previously described(*18*). Tumor growth was tracked via weekly caliper measurements. Once tumor growth became palpable, mice were randomized into groups with normalization of tumor size across groups, followed by the initiation of treatment. Animals were treated with 3 doses of IV carboplatin (8 mg/kg) or vehicle staggered by one day with 3 doses of anti-VEGFR3 antibody (100 μg) or IgG control as described above once tumors were palpable. Mice were euthanized by CO_2_ inhalation when largest tumors reached 2 cm in any direction; tumor-bearing and contralateral naïve fat pads were collected for histology as described above.

#### Studies in human xenograft breast cancer model

Human MDA-MB-231 breast cancer cells were cultured as described above. 1×10^6^ cells were suspended in 3.3 mg/ml growth-factor reduced matrigel in phosphate-buffered saline (PBS) and orthotopically injected in a subareolar fashion into the fourth mammary fat pad of immunocompromised female mice (NOD.CB17-Prkdcscid/J). Once tumors were palpable, carboplatin was administed in three injections as described above. Mice were euthanized by CO_2_ inhalation one week following final treatment; tumor-bearing and contralateral naïve fat pads were processed for histology as described above.

### Human tissue sample acquisition

Remnant, to be discarded, surgical resections (not needed for diagnostic purposes) of omental metastatic ovarian cancer and patient-matched normal omentum and benign pelvic mass omentum for immunohistochemical staining with podoplanin were collected into a tissue and data bank by waiver of consent and approved by the University of Virginia Institutional Review Board for Health Sciences Research. The UVA Biorepository and Tissue Research Facility procured remnant samples, including all breast cancer specimens and the majority of omental specimens, under this protocol from UVA Pathology for fixed and embedded specimens in paraffin. De-identified tissues and associated clinical data were pulled from this tissue bank and used in experiments approved by UVa IRB-HSR.

### Immunohistochemistry

Tumor-bearing mammary fat pads and lung tissues were dissected from mice and post-fixed in 4% PFA for 48 hours at 4°C; naïve fat pads underwent 24 hours of fixation. Fat pads were transferred to 70% ethanol for 24 hours, dehydrated, and paraffin-embedded. Tissues were sectioned at 7 μm thickness. Sections were stained with anti-podoplanin antibody (1 μg/ml, R&D Systems) followed by ImmPRESS HRP anti-goat IgG peroxidase/SG peroxidase detection (Vector Labs) and nuclear counter-staining with hematoxylin (Vector Labs) was performed. Human samples were processed and stained similarly. Slides were scanned at 20X on an Aperio Scanscope. For quantification of lymphatic vessel size, a custom interactive MATLAB (MathWorks) program utilizing the Image Processing Toolbox (MathWorks) was designed to identify and analyze lymphatic vessels. First, IHC images were binarized using an intensity threshold capable of isolating vessels with high specificity. Next, a flood-fill operation was used to uniformly fill the vessel area. These regions were then extracted for analysis. Vessels not captured or incompletely captured by the automated procedure were identified by user-drawn regions of interest (n=5/cohort, minimum of 20-30 representative vessels/cohort). Centroid coordinates, perimeter, and area was computed for each vessel. For lymphatic vessel density, all lymphatic (podoplanin+) vessels in the mammary fat pad were counted and vessel number was normalized to size of stromal area for each mouse to assess lymphatic vessel density as lymphatic vessel #/mm^2^ stroma. Intratumoral lymphatic vessels were rare and not included in these analyses in animal models. For lymphatic metastasis, sections were stained with anti-GFP antibody (5 μg/ml, Thermo Fisher, RFP Tag Monoclonal Antibody RF5R) and whole node confocal scans were used to quantify percent metastatic area of total node via image thresholding in ImageJ. Lungs were cryopreserved and embedded in OCT following fixation, cryosectioned, and stained with hematoxylin and eosin (H&E) to detect metastasis. Macroscopic metastatic lesions were counted by eye, whereas whole 20X scans of lung tissue were taken to detect micrometastatic lesions. Area of macroscopic lesions were measured in ImageJ.

### 3D *in vitro* co-culture model

10,000 LECs were seeded on the underside of 8 μm pore size 96-well tissue culture inserts (Corning). After 48 h, 50 μl of a Rat Tail Collagen I (Corning)/basement membrane extract (Trevigen) (0.18 mg/ml Collagen, 0.5 mg/ ml BME) containing Cell Tracker dye (ThermoFisher). Labeled human mammary fibroblasts (100,000 HMF/ml) and human breast cancer cells (660,000 TNBC cells/ml) was placed atop the inserts. After gelation, media was added to the bottom compartment and flow was applied via a pressure head in the top compartment overnight (~ 1 μm/s; 16-18 h), after which point docetaxel was applied via flow to the top compartment for 24 h and then flushed from the system with basal media. 48 h after drug application, gels were removed, dissociated using Liberase TM (Roche), and processed for flow cytometry. Inserts were processed for invasion analysis(*10*, *63*). All experimental conditions were run as triplicate samples in individual inserts.

### Invasion Assays

#### Tissue engineered 3D model of human breast tumor microenvironment

After gel was removed, tissue culture inserts were washed briefly in phosphate-buffered saline (PBS) and fixed with 4% paraformaldehyde. Inserts were stained with DAPI and visualized by fluorescence microscopy. Cancer cells (DAPI + Cell Tracker Deep Red+) were counted in five individual fields per well. Percent cancer cell invasion was calculated as previously described. Three technical replicates were averaged for each experimental run to give a single biological replicate value for statistical analysis.

#### Tissue homogenates from treated mice

Tissues from mice treated with 3 doses of carboplatin (as described above) were perfused with saline and dissected out. These tissues were degraded using 0.1 mg/ml Liberase TM for 30 minutes at 37C and then centrifuged down. The supernatant was removed and snap frozen until use. These samples were analyzed using BCA assay (Pierce) to determine total protein content. For invasion assays, 100,000 4T1 cells were resuspended in a 1.8 mg/ml collagen I (rat tail collagen, Corning), 0.2 mg/ml basement membrane extract (reduced growth factor, Trevigen) matrix which were loaded into a 96-well tissue culture insert plate (Corning). 25 μg of supernatants were placed in the lower chamber and cells invaded for 18h before gels were removed. Membranes were fixed and stained with DAPI and cells quantified to calculate total percent invasion based on total cells counted/total cells seeded.

### Flow Cytometry on *In Vivo* Tissues

Cells from *in vivo* digested lymph nodes and lungs were dissociated as previously described and stained with live/dead reactive dye, anti-mouse CD45 PerCP-Cy5.5 (eBioscience), anti-mouse CD31 FITC (eBioscience), and anti-mouse gp38 PE-Cy7 (eBioscience). Flow cytometry samples were processed using the Millipore Guava easyCyte 8HT Flow Cytometer and analyzed using InCyte software for total LEC counts per mg as well as percentage of LECs. Gating was done first on live cells, followed by CD45-populations. This subpopulation was gated for gp38 and CD31 with the following populations: CD31+gp38+ (LECs); CD31+gp38-(blood endothelial cells), and CD31-gp38+(fibroblastic reticular cells).

### Statistical Analysis

All data are presented as mean ± standard error of the mean (SEM). One-way or two-way ANOVA followed by Tukey’s multiple comparison test was used for statistical analysis of unmatched groups. Two group comparisons of normally distributed data as assessed by QQ plots were performed using unpaired t tests (with Welch’s correction if standard deviations were unequal), while comparisons of non-normal data were performed using Mann-Whitney U tests. Statistical analyses were run using Graphpad Prism software. *p<0.05* is considered statistically significant. All *in vitro* assays were performed with a minimum of three biological replicates unless otherwise noted, murine study numbers are noted in legends and by individual graphed data points. Graphs were generated using Graphpad Prism software and are shown with mean +/− standard error.

## Figure Generation

Figures were generated using Adobe Creative Suite (Photoshop and Illustrator). Schematics were generated using BioRender with license.

## Supplementary Materials

Table S1 – S6 Fig. S1 – S7

