## Supplemental Figures and Tables for "Platinum chemotherapy induces lymphangiogenesis in cancerous and healthy tissues that can be prevented with adjuvant anti-VEGFR3 therapy"

**Figure S1 | Lymphatic endothelial cell morphology in response to chemotherapies**

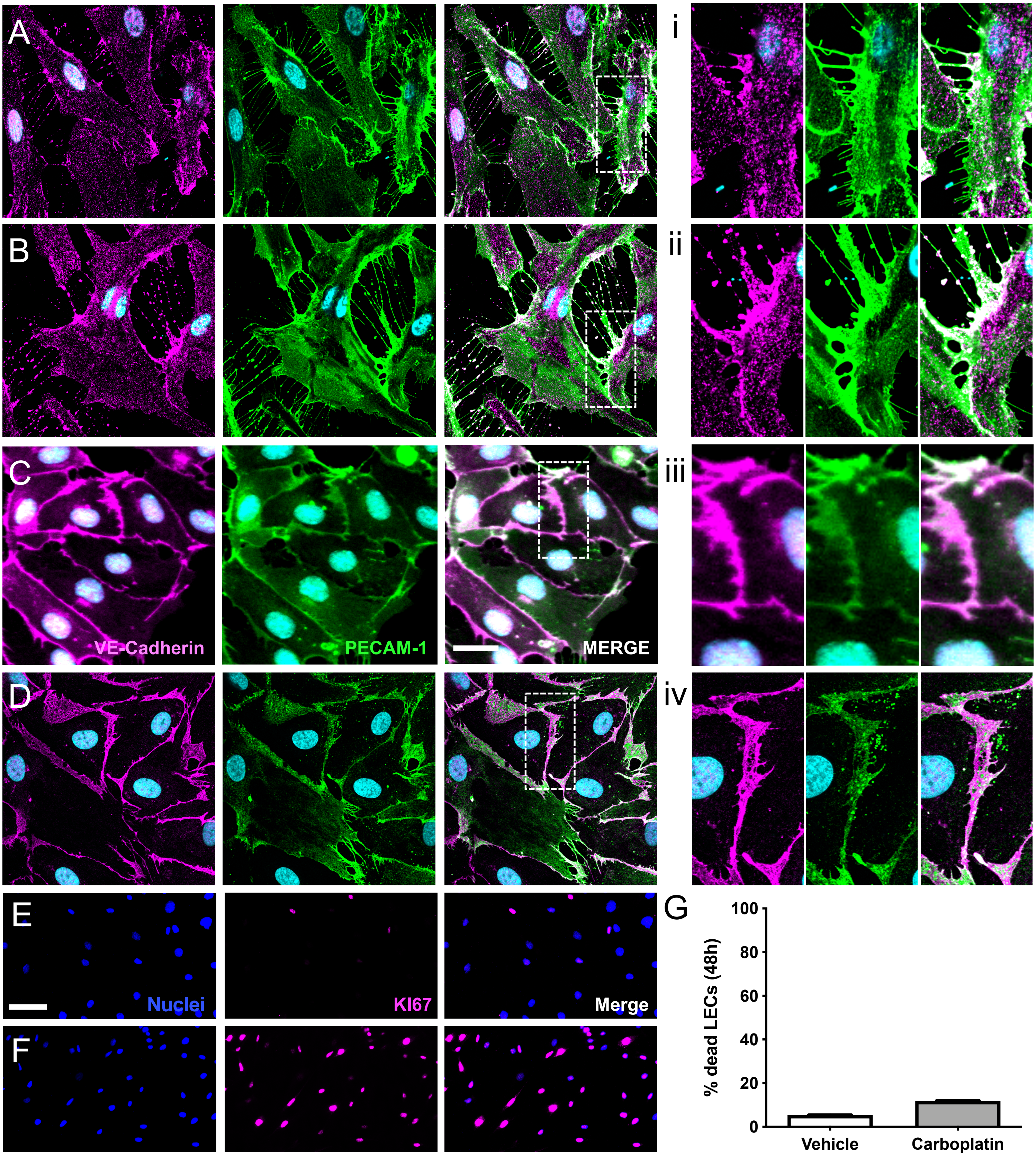


**Supplemental Figure 1 | Lymphatic endothelial cell morphology in response to chemotherapies. (A-D, i-iv)** Representative images of LEC morphology and junctions after platinum chemotherapy. Human LECs were treated *in vitro* with 1 µM of **(A)** cisplatin, **(B)** oxaliplatin, **(C)** doxorubicin, or **(D)** docetaxel for 6 hours. Cell morphology and junctions were visualized by immunocytochemical staining of VE-cadherin (magenta) and PECAM-1/CD31 (green) with DAPI (cyan) for nuclei. Junctions were imaged by high-powered confocal microscopy (scale bar = 25 microns). **(i-iv)** High magnification images of boxed areas in A-D. **(E-F)** Representative images of human LECs treated *in vitro* with **(E)** vehicle or **(F)** 1 µM of carboplatin and assessed for proliferation after 6 hours by immunocytochemical staining of KI67 (pink). (**G**) LEC viability after 48 hours of continuous treatment. Data represented as percentage of dead LECs as assessed by staining with live/dead viability dye, n=2.

**Figure S2 | Blood vasculature in a mesentery explant model**

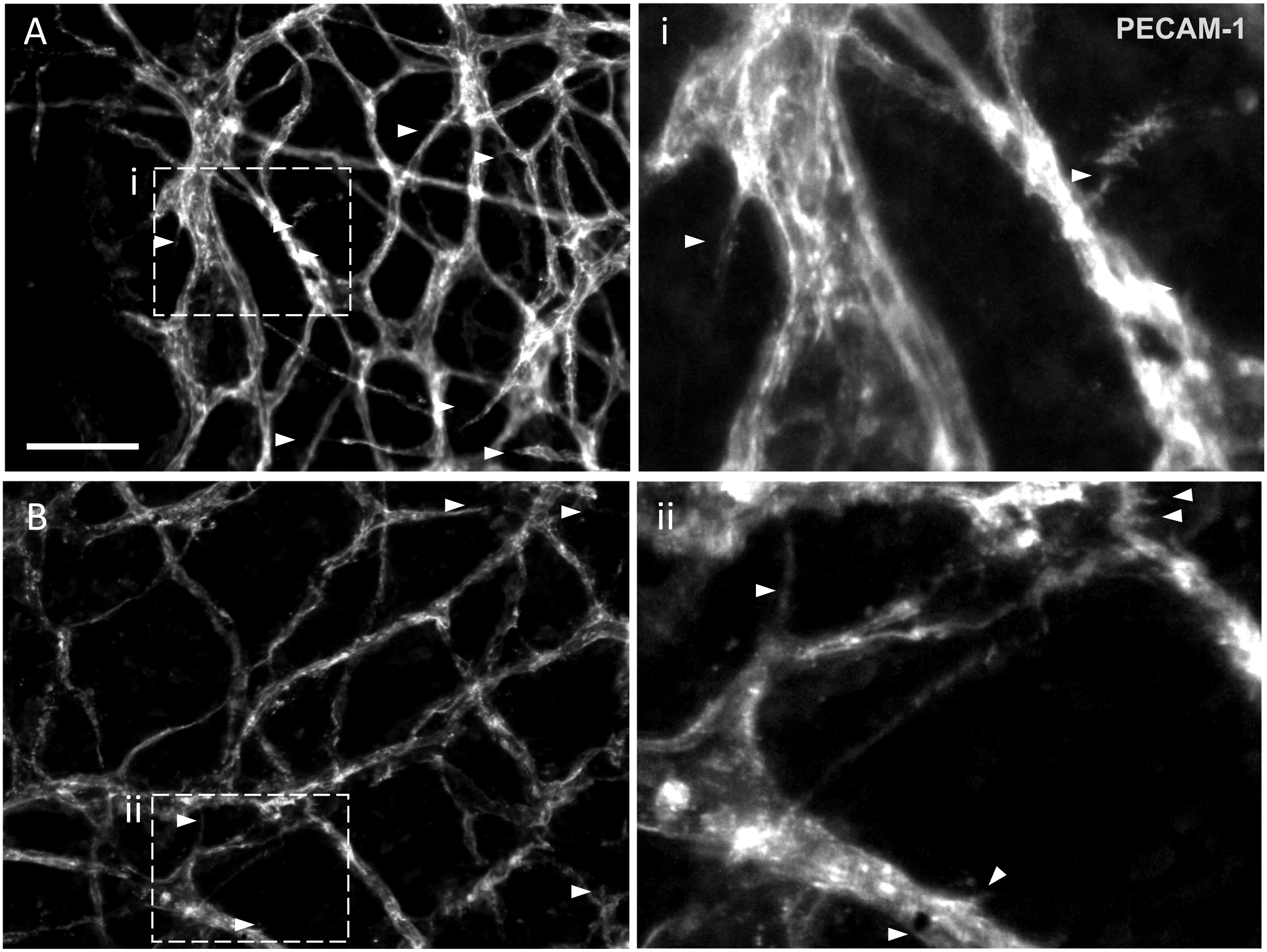


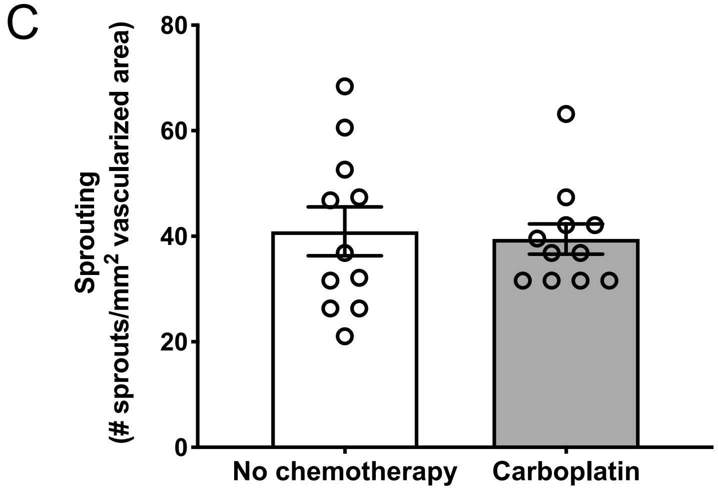


**Supplemental Figure 2 | Blood vasculature in the mesentery explant model. (A-C)** Representative images **(A, B)** and quantification **(C)** of cultured rat mesentery vascular networks treated with **(A)** vehicle or **(B)** 1 μM carboplatin for 5 days *ex vivo*. Vasculature was stained at endpoint with PECAM-1/CD31 (gray) for blood vessels. Arrowheads represent regions with high levels of sprouting. **(i, ii)** High magnification images of boxed areas in A and B. **(C)** Quantification is represented as number of sprouts per vascularized area. Scale bar=100 microns.

**Figure S3 | Multiple parameters of lymphatic vessels can be measured and quantified using specialized Matlab tool**

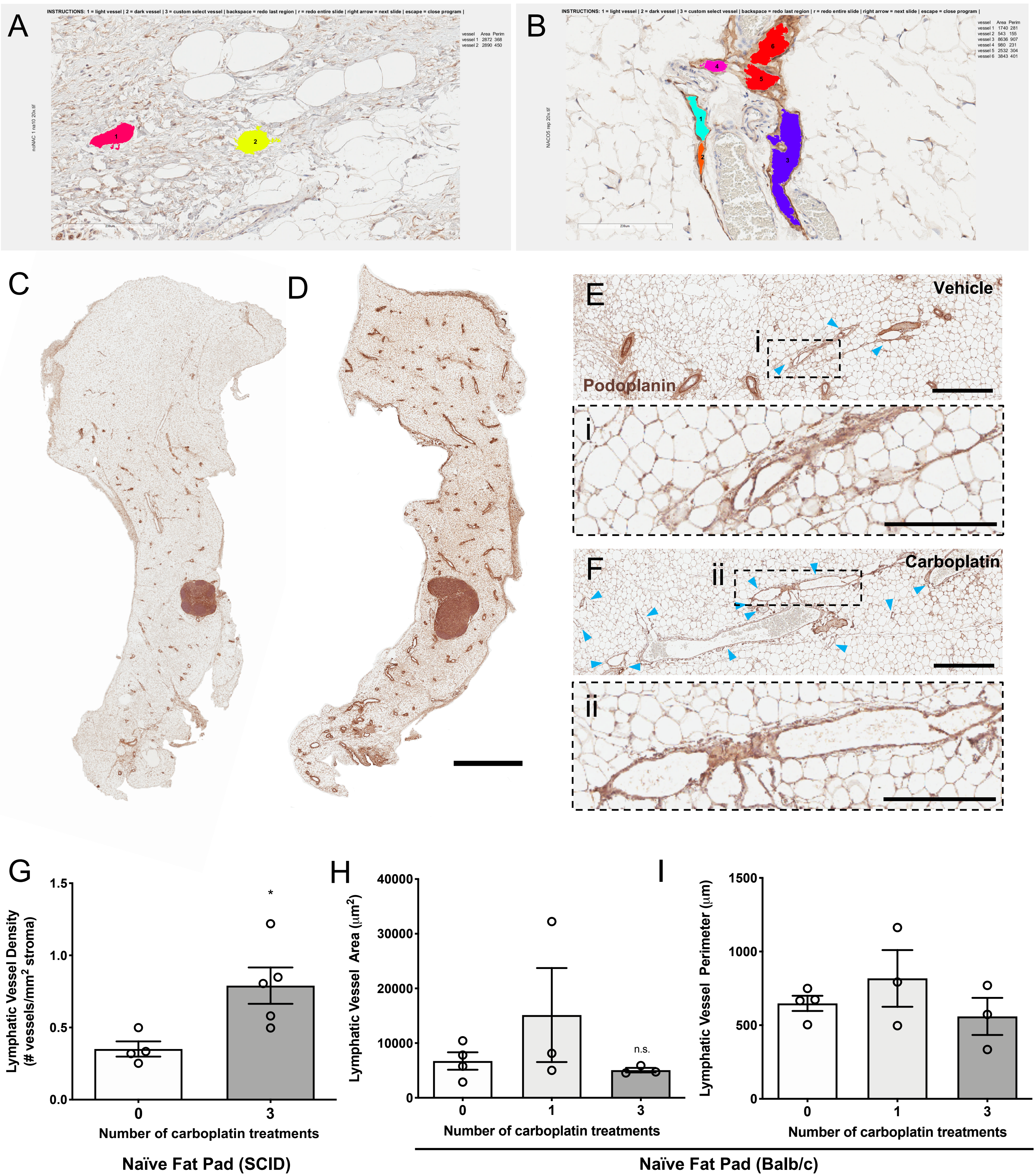

**Supplemental Figure 3 | Multiple parameters of lymphatic vessels can be measured and quantified using specialized Matlab tool. (A, B)** Representative images of lymphatic vessel quantification program using MathWorks Matlab tool. Colored and numbered regions are lymphatic vessels quantified by the software program. Vessel area and perimeter output can be seen on the far right of the window. **(A)** represents a sample with no chemotherapy while **(B)** represents a sample with chemotherapy. **(C-F, i-ii)** Representative images of mammary fat pads from naïve mice treated with 0, 1, or 3 doses of **(C)** vehicle or **(D)** 8 mg/kg systemic carboplatin by IV. Naïve fat pads were harvested three days following the final treatment and stained for lymphatic vessel marker podoplanin by immunohistochemistry. Representative images of lymphatic vessels are shown in **E** (vehicle) and **F** (carboplatin), with blue arrowheads showing lymphatic vessels (scale bar = 300 microns). (**i, ii**) High magnification images of lymphatic vessels from boxed regions (scale bar = 200 microns). **(G)** Immunocompromised NOD/SCID mice were injected with three doses of carboplatin (8 mg/kg, IV) or vehicle. Naïve mammary fat pads were harvested one week following final treatment and assessed for lymphatic vessel density by immunohistochemistry as previously described. Quantification of LVD in naïve mammary fat pad represented by total number of lymphatic vessels per stromal area. $^{*}p<0.05$ as analyzed by individual *t* tests, n=4-5 mice/cohort. **(H, I)** Quantification of lymphatic vessel (**H**) area and (**I**) perimeter in naïve balb/c MFPs treated as described in C-F.

### Figure S4 | Lymphatic analysis of 4T1 and xenograft mammary fat pads

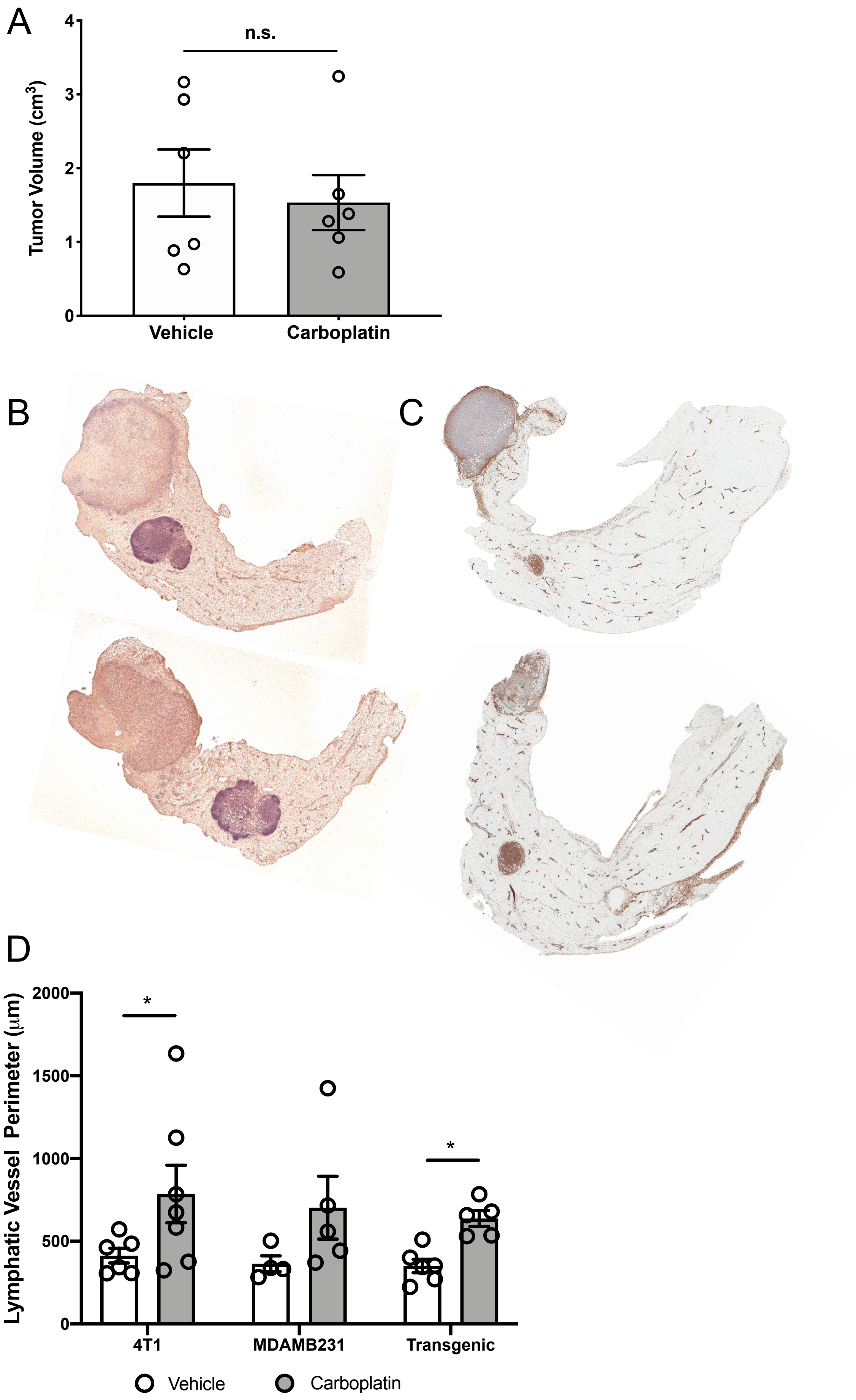


**Supplemental Figure 4 | Lymphatic analysis of 4T1 and xenograft mammary fat pads (A-D)** In all studies, mice were treated with carboplatin (8 mg/kg, 3 doses, IV) when tumors were palpable; mice were euthanized once the largest tumor within each study reached predetermined size endpoint. **(A)** Representative tumor weights at endpoint from autochthonous mammary tumor model, L-Stop-K-*KRas*$^{G12D}$*p53*$^{flx/flx}$-L-Stop-L-*Myristoylated p110α*-GFP$^{+}$ (induced by intraductal injection of adenovirus-Cre) show no difference in final tumor weight after treatment due to the low dose of carboplatin used. Representative tumor-bearing MFPs from **(B)** 4T1 (top, vehicle; bottom, carboplatin) and **(C)** xenografted MDAMB231 (top, vehicle; bottom, carboplatin) mouse models. **(D)** Quantification of lymphatic vessel perimeter from all tumor models treated as described in Fig. 1 and S1. *$p<0.05$ as analyzed by two-way ANOVA, n=4-6 mice/cohort.

**Figure S5 | Priming of tissues with carboplatin increases tumor cell invasion without altering tumor growth.**

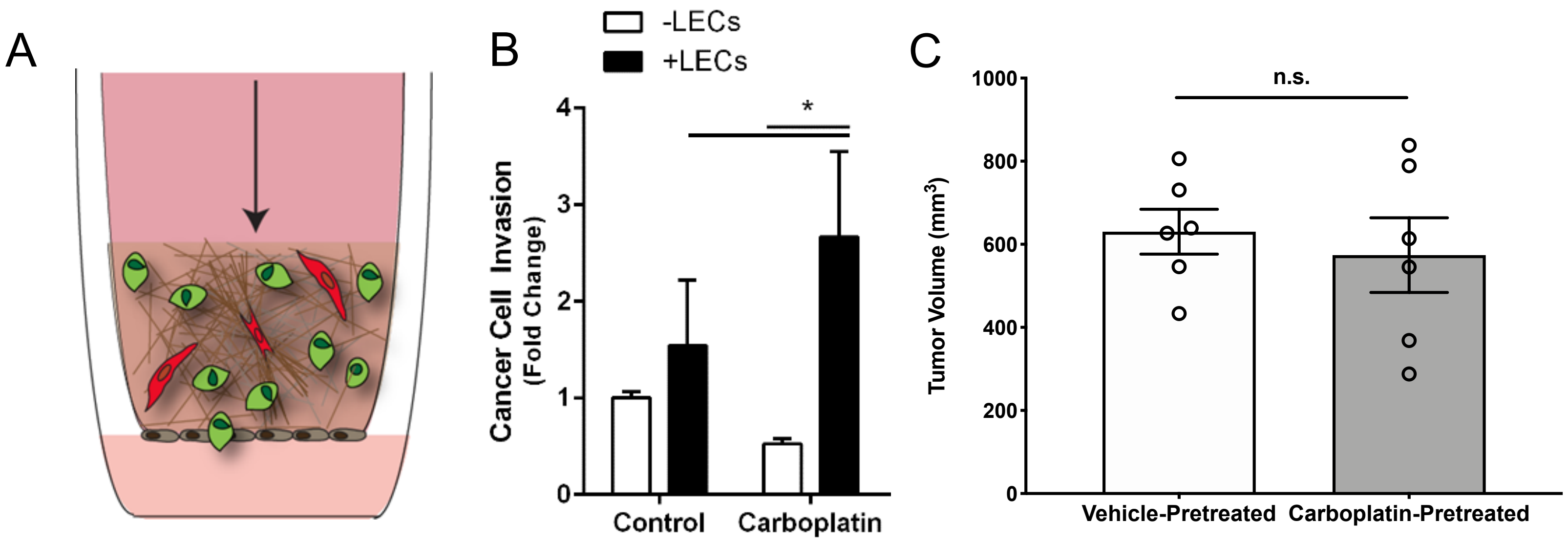


**Supplemental Figure 5 | Priming of tissues with carboplatin increases tumor cell invasion without altering tumor growth. (A)** Schematic of the *in vitro* tissue engineered model of the human breast cancer microenvironment including human triple-negative breast tumor cells, human lymphatic endothelial cells (LECs), and human mammary fibroblasts in a 3D collagen I matrix atop a porous tissue culture insert. Physiologically-relevant fluid flow with or without carboplatin is applied through the system via a pressure head. Schematic depicts experimental groups. **(B)** Fold change in invasion of MDA-MB-231 tumor cells across the porous membrane in our 3D microenvironment system +/- carboplatin treatment (1 μM) and/or in the presence or absence of LECs. *$p<0.05$. **(C)** Naïve mice were treated with carboplatin (8 mg/kg, 3 doses, IV) and drug was allowed to clear for one week after final treatment so as not to affect tumor initiation. Mice were then orthotopically implanted with 10,000 4T1 cells by subareolar injection. Tumors were allowed to grow until humane size endpoints were reached; there were no observed differences in final tumor size between cohorts.

**Figure S6 | Inhibition of VEGFR3 normalizes LECs *in vitro***

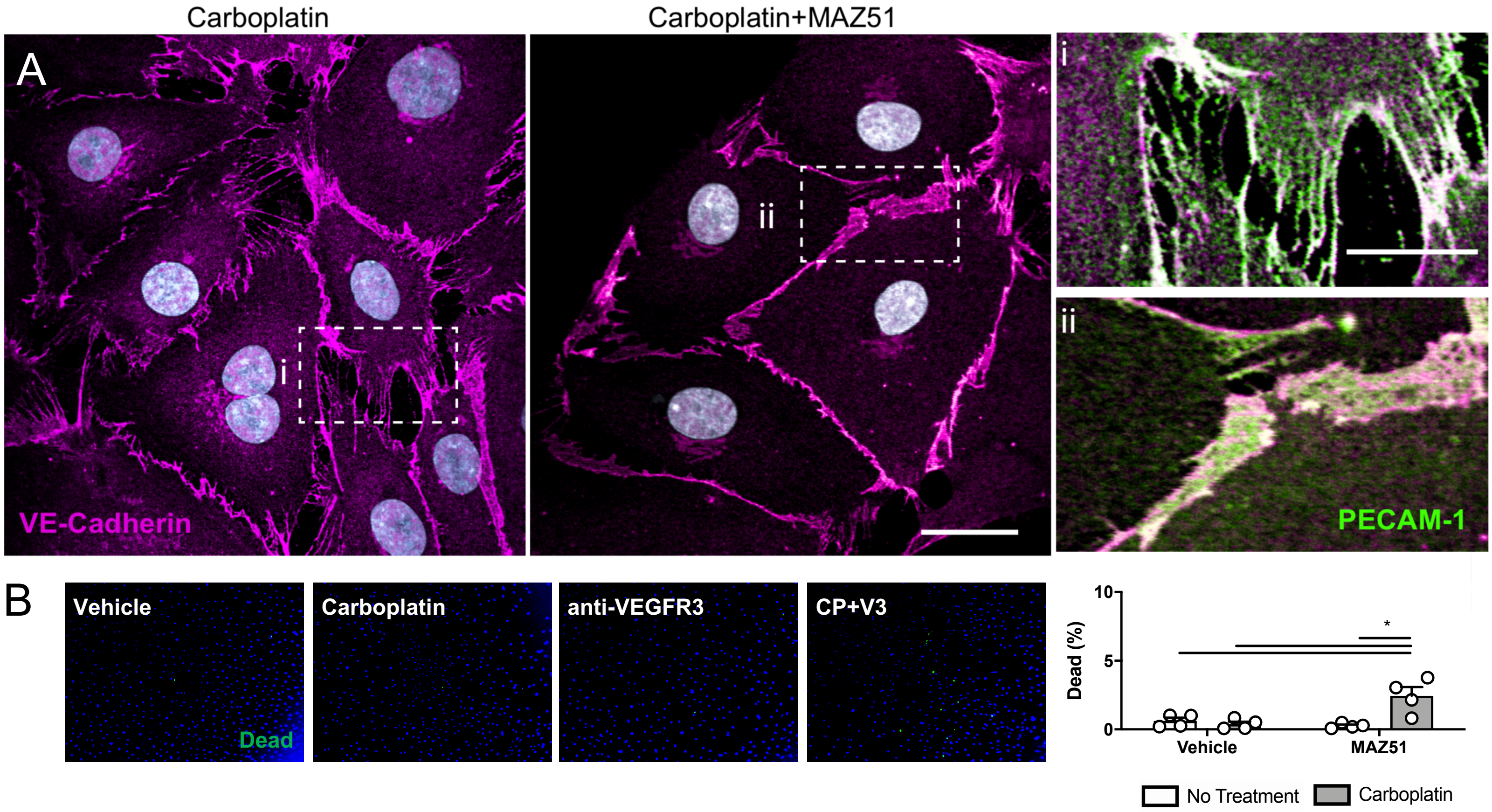


**Figure S6 | Inhibition of VEGFR3 normalizes LECs *in vitro*. (A)** LEC monolayers after 6h treatment VE-Cadherin (magenta) and PECAM1 (green) with high magnification images of boxed areas (i, ii). **(B)** Representative images and quantification of cell death as measured by percentage of cells dead per field after 48 hours of platinum agent (1μM) and/or MAZ51 and assessed by amine-based fluorescent reactive dye for dead cells. n≧3, *p<0.05

**Figure S7 | Inhibition of VEGFR3 mitigates all platinum-induced lymphatic changes**

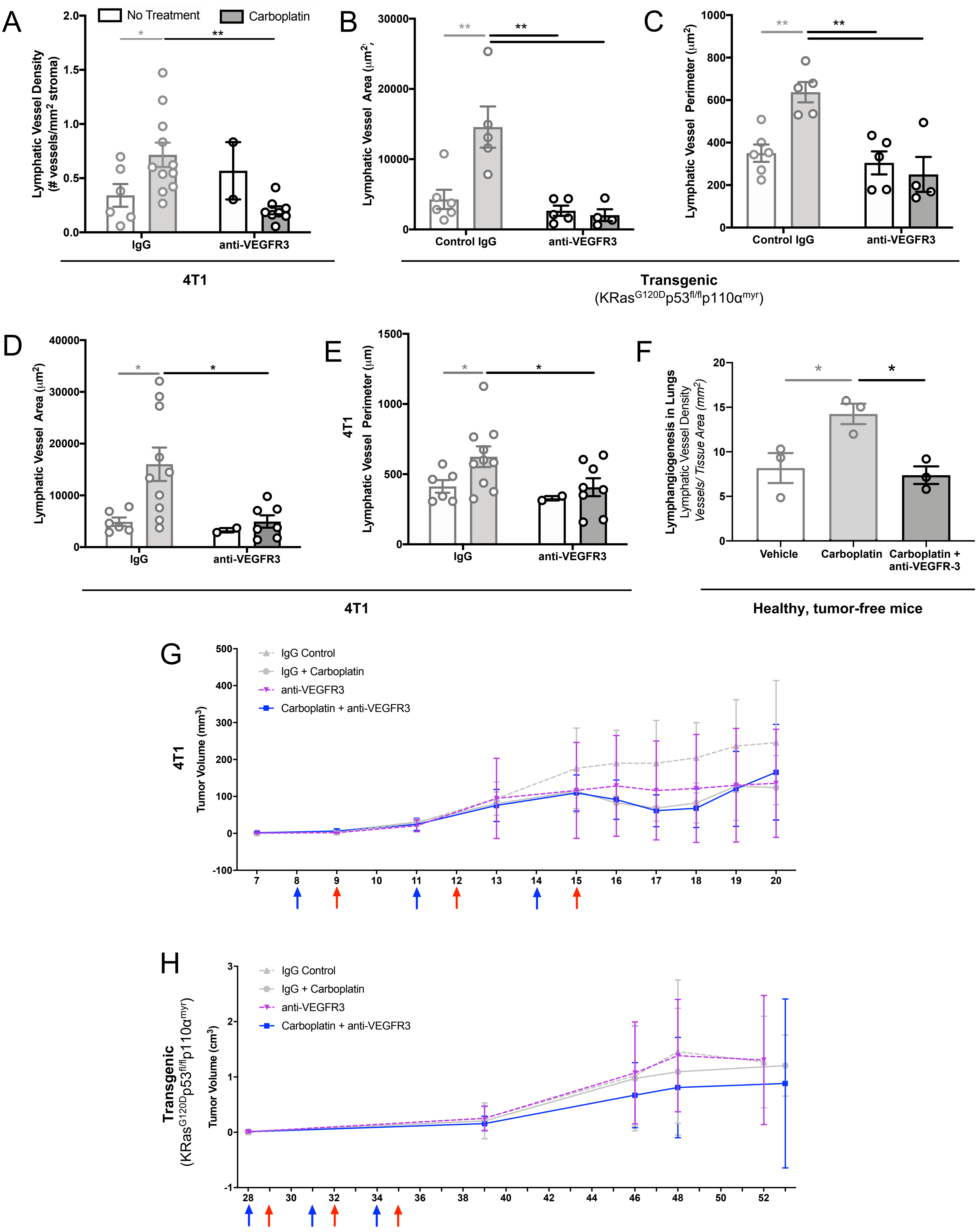

**Supplemental Figure 7 | Inhibition of VEGFR3 mitigates all platinum-induced lymphatic changes. (A)** Lymphatic vessel density as measured by number of podoplanin+ vessels per $mm^2$ stroma by immunohistochemical staining of tumor-bearing mammary fat pads from 4T1 mice treated with anti-VEGFR/control IgG antibodies + carboplatin/vehicle as previously described**. (B)** Lymphatic vessel area and (**C**) lymphatic vessel perimeter in tumor-bearing mammary fat pads of transgenic $KRas^{G120D}p53^{fl/fl}p110\alpha^{myr}$ mice treated with anti-VEGFR3/control IgG antibodies + carboplatin/vehicle as previously described. **(D)** Lymphatic vessel area and (**E**) lymphatic vessel perimeter in tumor-bearing mammary fat pads of 4T1 mice treated with anti-VEGFR3/control IgG antibodies + carboplatin/vehicle as previously described. **(F)** Naïve, healthy Prox1-tdTomato reporter mice were treated with carboplatin (8 mg/kg, 3 doses, IV) and lungs were collected 3 days following the final treatment. Lungs were fixed, sectioned, and assessed for lymphangiogenesis using image thresholding in Fiji. *$p<0.05$. **(G)** Tumor growth curve for 4T1 mice treated with anti-VEGFR3/control IgG antibodies + carboplatin/vehicle as previously described beginning once tumors were palpable. Blue arrows note antibody treatments, while red arrows note chemotherapy treatments. **(H)** Tumor growth curve for transgenic $KRas^{G120D}p53^{fl/fl}p110\alpha^{myr}$ mice treated with anti-VEGFR3/control IgG antibodies + carboplatin/vehicle as previously described beginning once tumors were palpable. Blue arrows note antibody treatments, while red arrows note chemotherapy treatments.

**Table S1 | Gene enrichment analysis of pathways upregulated in LECs after carboplatin treatment**

| Gene Enrichment Analysis by STRING | |
|---|---|
| **Pathways upregulated by carboplatin** | |
| *KEGG Pathway* | *Adjusted P-value* |
| MAPK signaling | 3.8E-05 |
| JAK-STAT signaling | 4.5E-05 |
| Pathways in cancer | 8.9E-05 |
| GAP junctions | 1.1E-04 |
| Cell adhesion molecules | 4.9E-04 |
| PI3K-AKT signaling | 1.0E-03 |
| HIF-1 signaling | 1.4E-03 |
| Ras signaling | 1.4E-03 |
| Chemokine signaling | 1.4E-03 |

**Table S3 | Intersecting pathways of microRNA regulation upregulated in LECs after carboplatin treatment.**

| MicroRNA Pathway Analysis by DIANA mirPath | | |
|---|---|---|
| **Intersecting pathways of miRNAs upregulated by carboplatin** | | |
| *KEGG Pathway* | *P-value* | *% of upregulated miRNAs involved* |
| TGFb signaling | 3.46E-08 | 33.3% |
| Adherens junctions | 6.21E-08 | 66.7% |
| Cell cycle | 0.006 | 44.4% |
| Proteoglycans in cancer | 0.014 | 66.7% |
| HIF-1 signaling | 0.019 | 44.4% |
| **Top miRNAs upregulated by carboplatin (FC>2)** | | |
| MIR3147, MIR130B, MIR3127, MIR4526, MIR4310, MIR421, MIR3116-1, MIR1224, MIR4659A | | |

**Table S2 | Pathways upregulated in LECs after carboplatin by RPPA analysis**

| RPPA Aanlysis | |
|---|---|
| **Pathways upregulated by carboplatin** | |
| *Signaling Pathway* | *Adjusted P-value* |
| Mammary gland development | 3.36E-05 |
| EGF/EGFR signaling | 6.96E-05 |
| Pac1-Pak1-p38-MMP2 | 7.46E-05 |
| FGF/ FGFR signaling | 8.39E-05 |
| ErbB2/ErbB3 signaling | 0.000379 |
| Pathways in cancer | 0.000651 |
| IL6 | 0.000995 |
| Pathways in TGF-B signaling | 0.00107 |
| HIF-1 signaling | 0.0019 |
| Alpha9 beta1 integrin | 0.00261 |
| Oxidative stress-induced senescence | 0.00292 |
| MicroRNAs in cancer | 0.00292 |
| Adherens junctions | 0.00329 |
| N-cadherin signaling events | 0.00402 |
| A6b1 and a6b4 integrin signaling | 0.00546 |
| Regulation of microtubule cytoskeleton | 0.00562 |
| Angiopoietin receptor Tie2-mediate signaling | 0.00596 |
| Posttranslational regulation of adherens junction stability and dissassembly | 0.00615 |
| Focal adhesion | 0.00615 |
| Cell-cell junction organization | 0.00748 |
| Beta1 integrin cell surface interactions | 0.00767 |
| Wnt signaling | 0.00797 |
| Alpha6 beta4 integrin | 0.00815 |
| DNA double strand break response | 0.00871 |
| MAPK signaling pathway | 0.012 |
| Cell cycle | 0.0137 |
| JAK/STAT signaling | 0.0138 |

**Table S4 | Patient cohort characteristics for primary breast tumor tissues from triple-negative breast cancer patients treated with and without neoadjuvant platinum chemotherapy.**

| Patient Characteristics (Tumor Tissues) | | | | | |
|---|---|---|---|---|---|
| Triple Negative Breast Cancer | | | | | |
| Patient ID | Clinical Staging | Tissue | Pathological Nodal Status | Neoadjuvant Chemotherapy | LVD (# LV/mm$^2$ tissue) |
| NA1 | Stage 1b | Mammary | pN0 | None | 0.12127603 |
| NA2 | Stage 1c | Mammary | pN0 | None | 0.25062657 |
| NA3 | Stage 1c | Mammary | pN0 | None | 0.22665947 |
| NA4 | Stage 1c | Mammary | pN0 | None | 0.66242999 |
| NA5 | Stage 2 | Mammary | pN0 | None | 0.28330206 |
| NA6 | Stage 2 | Mammary | pN0 | None | 0.39651071 |
| NA8 | Stage 2 | Mammary | pN0 | None | 0.80311657 |
| NA10 | Stage 2 | Mammary | pN0 | None | 0.19519879 |
| NA11 | Stage 1b | Mammary | pN0 | None | 0.18748535 |
| NA13 | Stage 1b | Mammary | pN0 | None | 0.4830745 |
| NA14 | Stage 1b | Mammary | pN0 | None | 0.26932668 |
| NA16 | Stage 1b | Mammary | pN0 | None | 0.30495553 |
| NA18 | Stage 1c | Mammary | pN0 | None | 0.45425667 |
| NA20 | Stage 2 | Mammary | pN0 | None | 0.5240027 |
| PA1 | Stage 1c | Mammary | pN1 | None | 0.5138908 |
| PA2 | Stage 2 | Mammary | pN1 | None | 0.32744592 |
| PA3 | Stage 1a | Mammary | pN1 | None | 0.12771678 |
| PA4 | Stage 1c | Mammary | pN1 | None | 0.2988741 |
| PA5 | Stage 3 | Mammary | pN1a | None | 0.22340023 |
| PA7 | Stage 2 | Mammary | pN1mi | None | 0.78431373 |
| PA8 | Stage 1a | Mammary | pN1a | None | 0.4642765 |
| NN4 | Stage 2 | Mammary | ypN0 | Cisplatin + Paclitaxel | 1.21918721 |
| PN5 | Stage 3a | Mammary | ypN3a | Cisplatin + Paclitaxel | 0.29317125 |
| PN6 | Stage 1a | Mammary | ypN1 | Cisplatin + Paclitaxel | 0.2788409 |
| NN5 | Stage 1c | Mammary | ypN0 | Cisplatin + Paclitaxel | 0.49746661 |
| PN3 | Stage 2 | Mammary | ypN1a | Cisplatin + Paclitaxel | 1.51354924 |
| PN11 | Stage 1c | Mammary | ypN1 | Carboplatin + Paclitaxel | 0.93373494 |

**Table S5 | Patient cohort characteristics for metastatic omentum from ovarian cancer patients treated with and without neoadjuvant platinum chemotherapy**

| Patient Characteristics (Tumor Tissues) | | | | | |
|---|---|---|---|---|---|
| Ovarian Cancer | | | | | |
| **Patient ID** | **Clinical Staging** | **Tissue** | **Pathological Diagnosis** | **Neoadjuvant Chemotherapy** | **LVD (# LV/mm$^2$ tissue)** |
| T0 | Stage 3 | Omentum | Positive | None | 0.089 |
| T3 | Stage 3 | Omentum | Positive | None | 0.807 |
| T4 | Stage 3 | Omentum | Positive | None | 0.186 |
| T5 | Stage 3 | Omentum | Positive | None | 0.119 |
| T6 | Stage 3 | Omentum | Positive | None | 0.419 |
| T8 | Stage 4 | Omentum | Positive | None | 0.427 |
| T9 | Stage 3 | Omentum | Positive | None | 0.254 |
| T10 | Stage 3 | Omentum | Positive | None | 0.566 |
| NACT0 | Stage 4 | Omentum | Positive | Carboplatin + Paclitaxel | 0.620 |
| NACT2 | Stage 3 | Omentum | Positive | Carboplatin + Paclitaxel | 4.272 |
| NACT3 | Stage 3 | Omentum | Positive | Carboplatin + Paclitaxel | 1.145 |
| NACT4 | Stage 3 | Omentum | Positive | Carboplatin + Paclitaxel | 2.906 |
| NACT5 | Stage 3 | Omentum | Positive | Carboplatin + Paclitaxel | 1.336 |
| NACT8 | Stage 3 | Omentum | Positive | Carboplatin + Paclitaxel | 0.219 |
| NACT9 | Stage 4 | Omentum | Positive | Carboplatin + Paclitaxel | 1.134 |
| NACT11 | Stage 3 | Omentum | Positive | Carboplatin + Paclitaxel | 0.563 |
| NACT12 | Stage 3 | Omentum | Positive | Carboplatin + Paclitaxel | 4.075 |

**Table S6 | Patient cohort characteristics for histologically benign/normal omentum from patients treated with and without platinum chemotherapy.**

| Patient Characteristics (Benign Tissues) | | | | |
|---|---|---|---|---|
| Histologically Normal Omentum | | | | |
| **Patient ID** | **Tissue** | **Pathological Diagnosis** | **Neoadjuvant Chemotherapy** | **LVD (# LV/mm$^2$ tissue)** |
| O1 | Omentum | Negative | None | 0.063 |
| O2 | Omentum | Negative | None | 0.560 |
| O4 | Omentum | Negative | None | 0.251 |
| NACO0 | Omentum | Negative | Carboplatin + Paclitaxel | 2.266 |
| NACO1 | Omentum | Negative | Carboplatin + Paclitaxel | 1.318 |
| NACO2 | Omentum | Negative | Carboplatin + Paclitaxel | 0.685 |
| NACO3 | Omentum | Negative | Carboplatin + Paclitaxel | 7.211 |
| NACO5 | Omentum | Negative | Carboplatin + Paclitaxel | 2.503 |
